# Naturally occurring variation in gene-associated transposable elements impacts gene expression and phenotypic diversity in woodland strawberry

**DOI:** 10.1101/2025.03.20.644342

**Authors:** Santiago Priego-Cubero, Rocio Tolley, Julia Llinares-Gómez, Camila Zlauvinen, Tuomas Toivainen, Timo Hytönen, Carmen Martín-Pizarro, Ileana Tossolini, Pablo A. Manavella

## Abstract

Transposable elements (TEs) are major components of plant genomes and powerful drivers of regulatory and phenotypic diversity, yet their mechanistic contributions to gene regulation in non-model crops remain poorly understood. Here, we generate a comprehensive and curated TE annotation for woodland strawberry (*Fragaria vesca*), revealing that TEs comprise ∼30% of the genome and uncovering an exceptional enrichment of gene-proximal miniature inverted-repeat transposable elements (MITEs) and short inverted repeats (IRs).

Integrating small RNA, DNA methylation and transcriptomic data across tissues and fruit developmental stages, we show that most gene-associated MITEs and IRs are epigenetically active, producing 24-nt siRNAs and displaying dynamic CHH methylation patterns during fruit ripening. Genome-wide analyses reveal that MITE- and IR-proximal genes occupy a distinct regulatory landscape compared to genes associated with other TE classes, consistent with a role in context-dependent transcriptional regulation.

As a mechanistic proof of concept, we demonstrate that a Mutator-like MITE located near *FvAAT1*, encoding a key alcohol acyltransferase required for aroma ester biosynthesis, modulates gene expression through siRNA-dependent DNA methylation and ripening-stage-specific chromatin reorganization. Transient perturbation of this MITE epigenetic state alters *FvAAT1* expression, and chromatin conformation capture reveals MITE-associated looping at this locus. Notably, this regulatory MITE is conserved in commercial strawberry species, suggesting an evolutionarily preserved function in fruit quality traits. Extending these findings to natural variation, we characterize thousands of TE insertion and deletion polymorphisms across 210 European *F. vesca* accessions and show that TE-based variation recapitulates population structure. A TE-informed genome-wide association analysis identifies significant associations between MITE and IR polymorphisms and fruit volatile compounds, including key aroma determinants.

Together, our results establish MITEs and short IRs as widespread, epigenetically active regulatory elements that shape gene expression, chromatin architecture and phenotypic diversity in strawberry. This work provides a conceptual and methodological framework to integrate TE-driven regulatory variation into population genomics, breeding and genome-editing strategies aimed at improving fruit quality.

## INTRODUCTION

Transposable elements (TEs) are repetitive, mobile DNA sequences widely dispersed across eukaryotic genomes. They constitute the most abundant genomic component in many species, accounting for nearly 60% of the human (*Homo sapiens*) genome [1] and up to 85% of the wheat (*Triticum aestivum*) genome [2]. Their extensive genomic presence directly impacts genome architecture, evolution, and gene regulation, particularly in response to environmental stressors [3–7].

TEs are classified into two major classes based on their transposition mechanisms [8]. Class I elements, or retrotransposons, transpose via an RNA intermediate through a “copy-and-paste” mechanism, whereas Class II elements, known as DNA transposons, move via a “cut-and-paste” mechanism [9, 10] or, in the case of Helitrons, a “peel-and-paste” replicative mechanism that uses a circular DNA intermediate [11]. TEs are further categorized into autonomous elements, which encode the proteins required for their own transposition, and non-autonomous elements, which rely on autonomous TEs for mobility [8].

Within these classes, TEs are subdivided into orders, superfamilies, and families based on structural and functional characteristics, such as replication strategies, sequence conservation, and enzymatic activity [8]. In plants, Class I Long terminal repeat (LTR) retrotransposons (e.g., Gypsy and Copia) are particularly abundant. Non-LTR retrotransposons (e.g., LINEs and SINEs), though more extensively studied in animals, also contribute significantly to plant genome evolution.

Beyond their contribution to genome size and structure, specific types of TEs are increasingly recognized for their regulatory potential. In this context, miniature inverted-repeat transposable elements (MITEs) represent a prominent group of short (<800 bp), non-autonomous Class II TEs that are highly abundant in eukaryotic genomes [7, 12, 13]. While many TEs accumulate in pericentromeric regions, MITEs and other short TE-derived inverted repeats (IRs) are frequently located near genes, where they can function as regulatory elements [14, 15]. A key feature that makes MITEs potent regulatory elements is their ability to form hairpin-shaped double-stranded RNAs (dsRNAs) upon transcription. These dsRNAs are processed by DCL3 into 24-nt small interfering RNAs (siRNAs) that trigger DNA methylation through a RNA-dependent RNA Polymerase 2 (RDR2)-independent variant of the RNA-directed DNA methylation (RdDM) pathway. This non-canonical form of RdDM plays a crucial role in silencing TEs to maintain genome stability [16–18]. Different from the canonical RdDM-initiated silencing of TEs through heterochromatin formation, RdDM-mediated DNA methylation of MITEs does not commonly act as a silencing mark but instead could impact the expression, both positively and negatively, of nearby genes [7].

TE research has expanded significantly with the increasing availability of genome sequences, enabling improved annotation, classification, and functional analysis of these elements. However, challenges remain, particularly in non-model species with incomplete TE libraries [19]. Various computational approaches have been developed for TE annotation [20], including *de novo* prediction [21–23], motif-based searches [24, 25], and similarity-based methods [26, 27]. Combining multiple strategies has proven most effective in generating accurate and comprehensive TE annotations [19, 23, 28, 29].

One of the most compelling open questions in TE research is understanding how the large natural variation in TE content within a species contributes to phenotypic diversity. Thanks to advancements in long-read high-throughput sequencing technologies, the study of TE polymorphisms has gained attention due to their role in complex traits and their potential as genetic markers for plant research and breeding [30, 31]. Integrating TE polymorphisms with Genome-Wide Association Studies (GWAS) has provided valuable insights into the functional implications of TEs in various biological processes [32–35]. For instance, transposon insertion polymorphisms (TIPs) have been shown to improve predictions of agronomic traits beyond what is possible with single-nucleotide polymorphism (SNP)-based approaches alone [34, 36–38].

The woodland strawberry (*Fragaria vesca*) is a wild diploid species from the Rosaceae family that serves as a model for strawberry research. It is one of the four ancestral species contributing to the polyploidization of the cultivated octoploid *F. × ananassa*, the most widely consumed strawberry species. Among the four subgenomes of *F. × ananassa*, the *F. vesca* subgenome exhibits the lowest density of TEs near genes and encodes significantly more dominantly expressed homeologs than the other three subgenomes combined [39]. While previous studies have examined TE diversity in the *F. vesca* genome [40–42], their complete annotation, functional characterization, and linkage with phenotypic variation remain poorly explored.

In addition to its genomic significance, *F. vesca* fruits are distinguished by their unique volatilome, which contributes to their intense floral and fruity aroma [10, 43]. The species exhibits substantial phenotypic diversity influenced by both genetic variation and environmental factors. The contribution of TEs to this phenotypic diversity remains largely unexplored. Given its importance as a genetic resource, understanding TE-driven variation in *F. vesca* could provide valuable insights into fruit development and quality traits, with potential applications for breeding.

In this study, we systematically annotated transposable elements in the *Fragaria vesca* reference genome, generating a curated TE library. Using this resource, we analyzed TE insertion and deletion polymorphisms across 210 re-sequenced European *F. vesca* accessions [44], revealing their contribution to genetic diversity and subpopulation structure. We then integrated TE polymorphisms into a genome-wide association framework, identifying significant associations with economically important fruit traits, including volatile compound profiles.

As a proof of concept, we functionally validated the regulatory impact of a MITE located near FvH4_7g18570, which encodes the alcohol acyl transferase *FvAAT1*, a putative ortholog of *SAAT* involved in volatile ester biosynthesis in *F. × ananassa [45]*. Combining gene expression analyses with chromatin conformation profiling at this locus, we demonstrate that MITE-derived siRNAs trigger DNA methylation that reshapes local chromatin organization, thereby modulating *FvAAT1* expression in a ripening-stage-dependent manner.

While genome-wide TE distribution patterns, such as LTR enrichment in pericentromeric regions and MITE localization near genes, have been described in other plant species, the TE landscape of *F. vesca* has remained poorly annotated and functionally unexplored. Our work provides a comprehensive TE resource for this species and establishes a framework to investigate the regulatory potential of MITEs and short inverted repeats, particularly during fruit development. By integrating TE-driven variation into population-genomic and functional analyses, this study offers new insights into how TEs contribute to gene regulation and phenotypic diversity in strawberry, and highlights their potential as targets for breeding and genome-editing strategies aimed at improving fruit quality.

## RESULTS

### Comprehensive identification and annotation of transposable elements in *Fragaria vesca*

Transposable elements (TEs) do not typically experience selective pressures that preserve their sequences over time. Consequently, they tend to become fragmented due to the accumulation of random mutations, which drives divergence among different TEs and complicates their identification [23, 46, 47]. To address these challenges, we combined two recent bioinformatic pipelines, EDTA [48] and DeepTE [49], and applied TEtrimmer [50], a consensus refinement and de-redundancy tool, to generate a semi-curated de novo TE library in *F. vesca* (Figure S1). This allowed us to comprehensively identify and classify TEs in the *F. vesca* v4.0 reference genome [51, 52] into classes, orders, and superfamilies (Table 1, Figure 1A).

**Figure 1.**
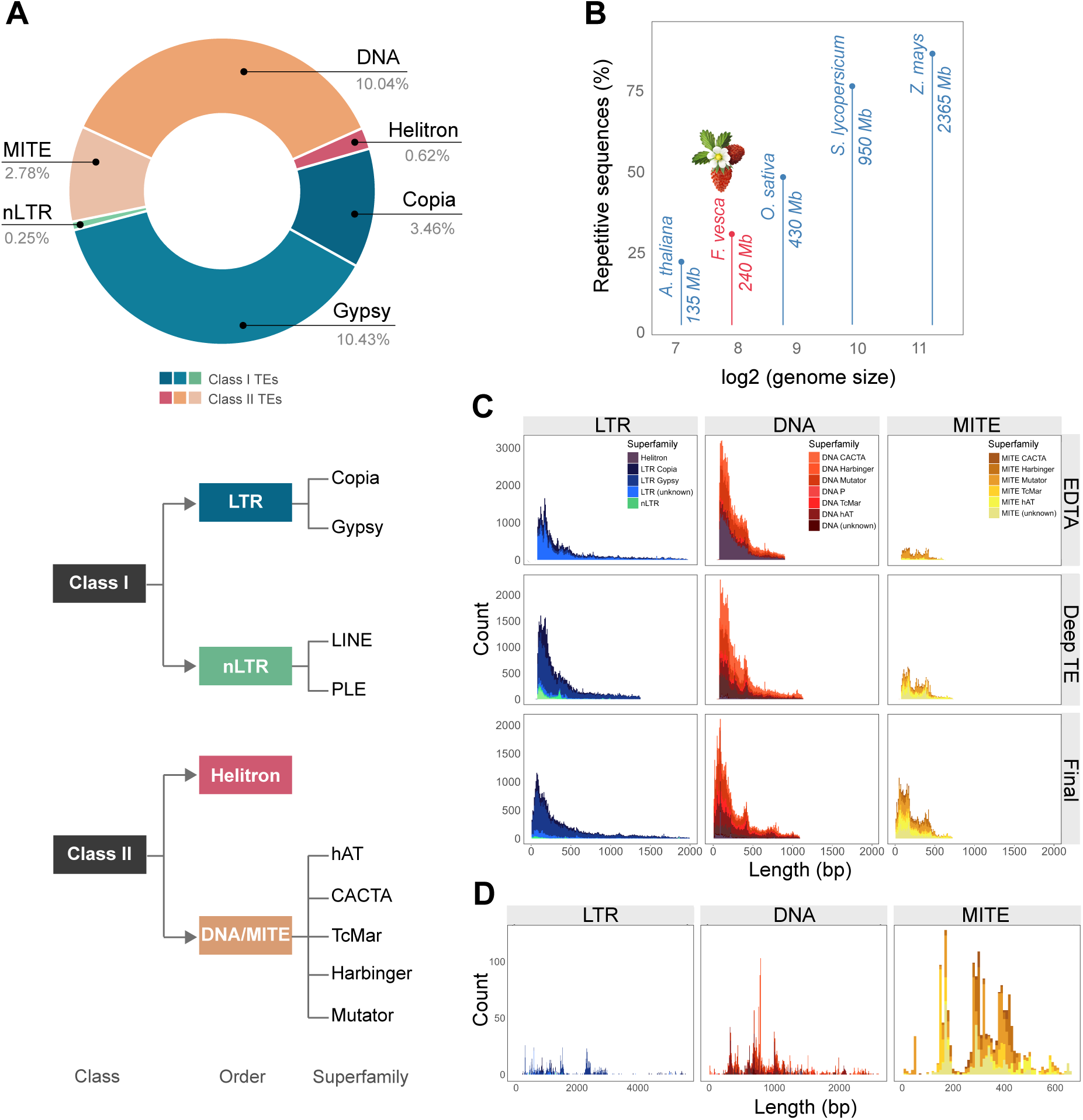
Transposable Element (TE) diversity in *Fragaria vesca*. **(A)** Total coverage of the *F. vesca* genome assembly by the main TE classes, shown as percentages relative to the entire genome. Representation of TE classification scheme generated in this study is displayed. **(B)** Repetitive sequence content as a proportion of total genome size in several plant species, listed in ascending order: *Arabidopsis thaliana*, woodland strawberry (*Fragaria vesca*), rice (*Oryza sativa*), tomato (*Solanum lycopersicum*), and maize (*Zea mays*). **(C)** Length distribution of TEs (including fragmented TEs) in the *F. vesca* v4.0.a2 annotation. The classifications are listed in descending order for EDTA, DeepTE, and the curated annotation. **(D)** Length distribution of intact TEs in the curated *F. vesca* v4 annotation.

**Table 1.** Number of unique TEs, number of copies, and total TE coverage of the *F. vesca* genome assembly v4.0, according to their classification.

| Classification | N° of TEs | N° of Copies | Genome Coverage (%) |
| --- | --- | --- | --- |
| **Class I** |  |  |  |
| **LTR** |  |  |  |
| Copia | 131 | 8,647 | 3.46 |
| Gypsy | 262 | 25,977 | 10.43 |
| Unclassified | 35 | 3,536 | 0.91 |
| Total LTRs | 428 | 38,160 | 14.80 |
| **non-LTR** |  |  |  |
| LINE (L1) | 8 | 292 | 0.07 |
| PLE | 7 | 548 | 0.18 |
| Total non-LTRs | 15 | 840 | 0.25 |
| **Total Class I** | **443** | **39,000** | **15.05** |
| **Class II** |  |  |  |
| **Subclass 1** |  |  |  |
| **MITE** |  |  |  |
| CACTA | 7 | 1,084 | 0.08 |
| Harbinger | 25 | 4,243 | 0.46 |
| Mutator | 56 | 7,495 | 0.77 |
| TcMar | 23 | 2,721 | 0.28 |
| hAT | 53 | 5,780 | 0.68 |
| Unclassified | 28 | 4,737 | 0.51 |
| Total MITEs | 192 | 26,060 | 2.78 |
| **DNA (non-MITE)** |  |  |  |
| CACTA | 50 | 8,495 | 2.97 |
| Harbinger | 20 | 2,376 | 0.49 |
| Mutator | 166 | 13,946 | 1.95 |
| TcMar | 63 | 7,391 | 1.66 |
| hAT | 134 | 12,237 | 2.00 |
| Unclassified | 63 | 6,184 | 0.96 |
| Total DNA | 496 | 50,629 | 10.03 |
| **Subclass 2** |  |  |  |
| Helitron | 13 | 2,291 | 0.62 |
| **Total Class II** | **701** | **78,981** | **13.43** |

The complete set of identified TEs accounted for approximately 30% of the *F. vesca* genome (Figure 1B). In plants, TE content is typically proportional to genome size [53, 54]. For example, the maize genome (∼2,500 Mb) contains over 80% TEs and repetitive elements, whereas the smaller *Arabidopsis thaliana* genome (∼135 Mb) contains only 21% TEs [55–57]. The proportion of TEs identified for *F. vesca* (∼30%), which has a genome size of ∼220 Mb [58], aligns with these observations (Figure 1B).

In contrast to most plant species, where Class I retrotransposons typically dominate, the genome of *F. vesca* exhibits a relatively balanced proportion between Class I and Class II elements. Class I retrotransposons constitute approximately 15% of the total genomic DNA, whereas Class II DNA transposons represent around 13.4% (Table 1). However, domain-based analyses revealed that only a small fraction of Class II elements retain detectable transposase-associated domains, indicating that most DNA transposon copies correspond to non-autonomous or fragmented elements rather than potentially autonomous transposons. Within Class I, LTR retrotransposons remain the most prevalent, comprising most of these elements, with the Gypsy superfamily being particularly abundant (Figure 1A), and non-LTR retrotransposons contribute only a minor fraction (∼0.25%) of the genome. The relatively moderate proportion of Class I retrotransposons observed in *F. vesca* might reflect its smaller genome size compared to other plant genomes and could be further influenced by limitations in accurately assembling centromeric regions, which are enriched with these elements.

Interestingly, *Fragaria* species exhibit higher diversity among DNA transposons, compared to other plant genomes, highlighting the unique composition of their genomes [40]. After re-classifying DNA transposons into MITES and non-MITEs with DeepTE, almost 13% of identified TEs belong to subclass 1, including both MITEs and other DNA transposons from the CACTA, Harbinger, TcMar, hAT, and Mutator superfamilies. The remaining 0.62% corresponds to subclass 2, specifically Helitrons (Figure 1A). Despite their known difficulty in identification, the proportion of Helitrons identified in the *F. vesca* genome is consistent with reports in other plant genomes, where they typically constitute below 4% of the genome [40, 48, 59].

We then analyzed the size distribution of TEs for each class based on the annotation generated for *F. vesca*, including both fragmented and intact TEs (Figure 1C, Figure S2, Table S1). Overall, no clear differences were observed in size distributions among the results obtained from EDTA, DeepTE, and the curated library (Figure 1C). The average length of each intact and fragmented TE type is detailed in Figure S2 and Table S1. The similarity in size distributions across libraries suggests that the final curated TE library preserves the structural characteristics of both intact and fragmented elements, supporting the robustness of the library for subsequent analyses.

A high proportion of fragmented TEs was identified across the genome. This explains the relatively low mean length values observed for the complete TE library compared to the intact subset (Figure S2), a pattern consistent with the long-term accumulation of mutations and deletions in ancient TEs. While Class I elements generally have lengths below 4 kb, the longest sequences belong to the LTR order (Figure 1D, Figure S2). In contrast, Class II TEs display a broader size range strongly linked to their respective superfamily classification (Figure 1D). The distinct size patterns observed among TE classes suggest differences in replication strategies and selective pressures acting on different TE families.

To better understand the structural preservation of these elements, we partitioned the library into two mutually exclusive sets: intact and fragmented elements. A marked contrast in the distribution of TE classes between these groups was observed. While DNA transposons dominated both fractions, they showed a significant increase of 12.3% in the intact library compared to the fragmented set. This enrichment was driven primarily by MITEs, which accounted for 36.6% of the intact elements, a substantial increase of 16.1% relative to their 20.5% representation in the fragmented library. Conversely, LTRs were more prevalent among fragmented sequences than in the intact subset, representing a decrease of 9.5%. This highlights a higher rate of structural decay for these larger elements. The abundance of intact MITEs, which are known for being relatively short, can be attributed to their unique structural nature. Unlike autonomous DNA transposons or LTR retrotransposons, MITEs are characterized by their compact sequence length (Figure 1D) and lack of internal coding regions. From a genomic perspective, a shorter sequence presents a significantly smaller “mutational target” for stochastic fragmentation, deletions, or nested TE insertions. While the integrity of an LTR element is easily compromised by a single internal deletion, MITEs are more likely to remain structurally recognizable as “intact” over evolutionary time. Furthermore, because MITEs do not rely on internal Open Reading Frames (ORFs) for their identification they are less sensitive to the sequence decay that typically fragments autonomous elements. Consequently, while larger autonomous elements are rapidly converted into genomic “noise” or fragmented relics, MITEs maintain higher structural stability.

### Genome-wide distribution and abundance of Transposable Elements

The distribution, abundance, and genome coverage of TEs vary substantially across species and are strongly influenced by subfamily classification. While some TEs tend to accumulate near protein-coding genes or toward the centromeres, others are more evenly distributed across chromosomes. To assess the distribution of TE types in *F. vesca*, circos ideograms were generated for each TE class, with gene density included as a reference (Figure 2).

**Figure 2.**
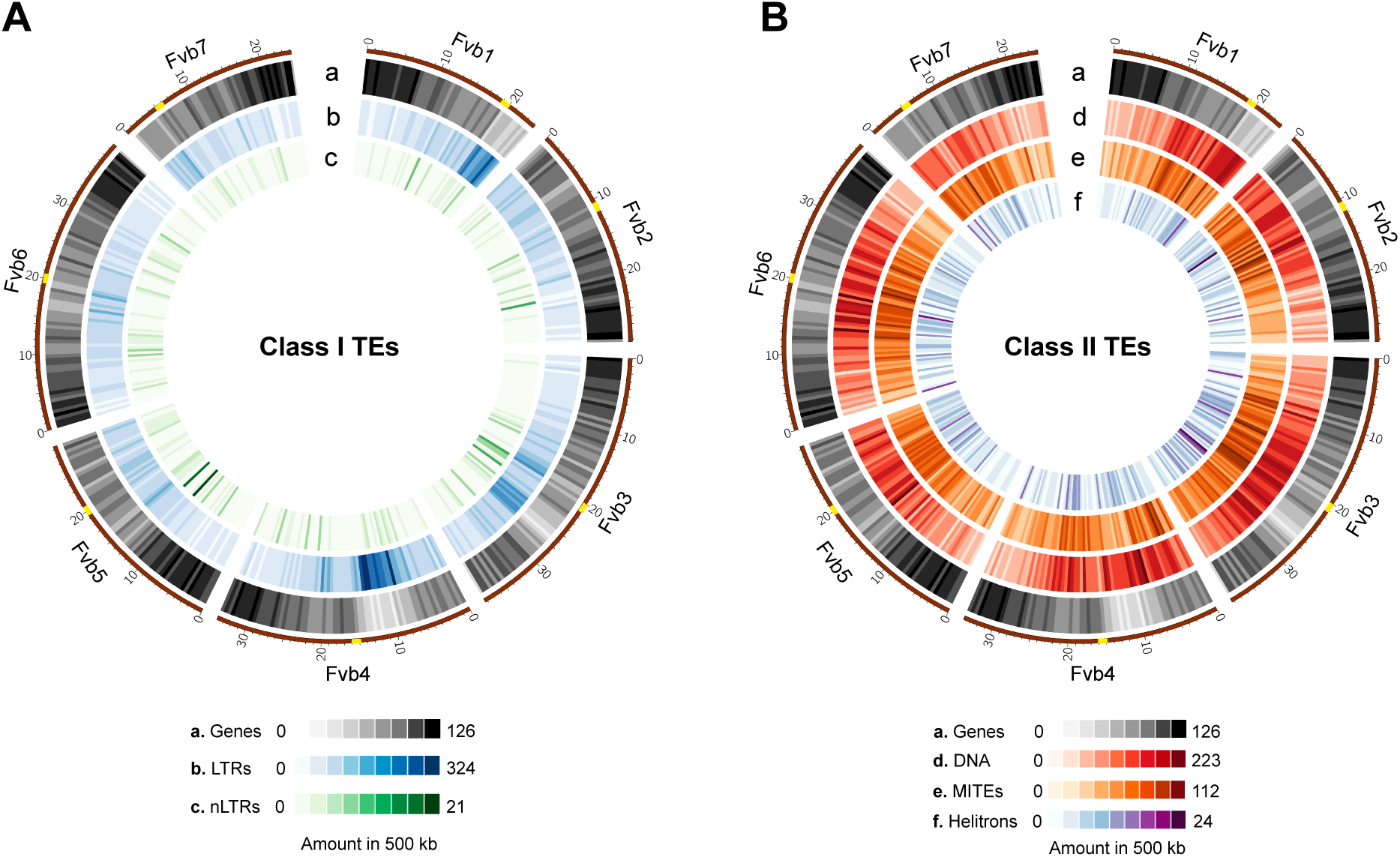
Distribution of TEs in the *F. vesca* genome, represented as density plots within 500 kb windows. **(A)** Class I TEs: LTR (Track b in blue), non-LTR (Track c in green). **(B)** Class II TEs: DNA (Track “d” red), MITE (Track “e” in orange), Helitron (Track “f” in purple). The color scale indicates the number of elements found within each 500 kb window. The distribution of genes is shown in grayscale (Track “a”) in both graphs. Centromeres are marked as a yellow dot in the brown outer line.

As shown in Figure 2A, regions with higher Class I TE content tend to have lower gene density, a pattern particularly evident for chromosomes Fvb1, Fvb3, and Fvb4. This TE distribution is predominantly driven by LTR transposons, which are the most abundant among Class I. A large fraction of LTR elements appeared located more than 3,000 bp from genes, suggesting a low likelihood of gene association (Figure S3). This distribution pattern of Class I elements has been previously reported in several plant species, including tomato (*Solanum lycopersicum*) and rice (*Oryza sativa) [56, 57, 60]*).

In contrast, Class II TEs are more concentrated near pericentromeric regions compared to chromosome arms except for Helitrons, which exhibit a more widespread distribution across the genome (Figure 2B). Helitrons positioning varies considerably between species. For example, Helitrons display a more erratic pattern of distribution in potato and rice [57, 61] compared to Arabidopsis, where they are enriched in gene-poor pericentromeric regions, or in maize (*Zea mays*), where they are more abundant in gene-rich regions [62].

MITEs exhibit a distinct distribution pattern characterized by a notable concentration in gene-rich regions, particularly in the immediate vicinity of, or within, coding sequences (Figure S3). This pattern aligns with previous reports describing the regulatory role of MITEs in gene expression [56, 57, 63]. Notably, 46% of *F. vesca* genes harbor a MITE within 3,000 bp of their coding sequence, and 15% of these genes contain MITEs within their coding regions. This proximity to genes reinforces the potential role of MITEs in influencing gene expression through epigenetic and chromatin-level mechanisms encouraging us to focus on the study of these elements.

### Characterization of MITEs and short TE-derived IRs in *F. vesca*

MITEs are frequently found in close proximity to genes, where they influence gene expression and act as evolutionary agents contributing to adaptation and developmental processes [64, 65]. This regulatory impact is often mediated by the production of 24-nt small RNAs through the non-canonical RdDM pathway [64, 66, 67]. Such processes trigger localized DNA methylation and induce structural changes in chromatin organization, leading to the formation of short-range chromatin interactions that modulate the transcriptional activity at neighboring genes [64–67]. A similar effect can be triggered by gene-associated TE-derived short inverted repeats (IRs) elements, which often originate from MITEs that have lost distinctive structural elements. The presence of IRs reflects the progressive fragmentation and degeneration of TEs over evolutionary time [7]. In any case, and for the purpose of the regulatory effects studied here, IRs and MITEs operate similarly.

Given the mechanistic similarities between IRs and MITEs, we employed *einverted* from the European Molecular Biology Open Software Suite (EMBOSS) [68] to also annotate all IRs in *F. vesca*. We then classified the putative nature of the identified IRs and compared them with the superfamilies previously assigned to MITEs (Figure 3A). IRs can originate from both MITE-type and DNA transposons, Class I elements, or even non-TE sequences [64]. Consistently, 84% of the identified IRs were classified as MITE-derived elements, in line with the matching distribution patterns of MITEs and IRs (Figure S4A). Additionally, the overall superfamily distribution of IRs was consistent with that of the annotated MITEs in our TE library (Figure 3A). However, the proportion of IRs from the MITE Harbinger superfamily was lower than that of the annotated MITEs. This suggests that certain IRs may have originated from more ancient MITE elements that have since diverged from their original sequence patterns. Lastly, 16% of the IRs were classified as elements distinct from MITEs (“Other” category in Figure 3C), predominantly belonging to DNA Mutator/CACTA, LTR Gypsy/Copia, and unknown origin (Figure S4B). Considering this classification, we observed that 58% of the identified IRs overlapped by at least 20% of their length with MITEs annotated in our TE library. Therefore, subsequent analyses were performed separately for MITEs and MITE-independent IRs.

**Figure 3.**
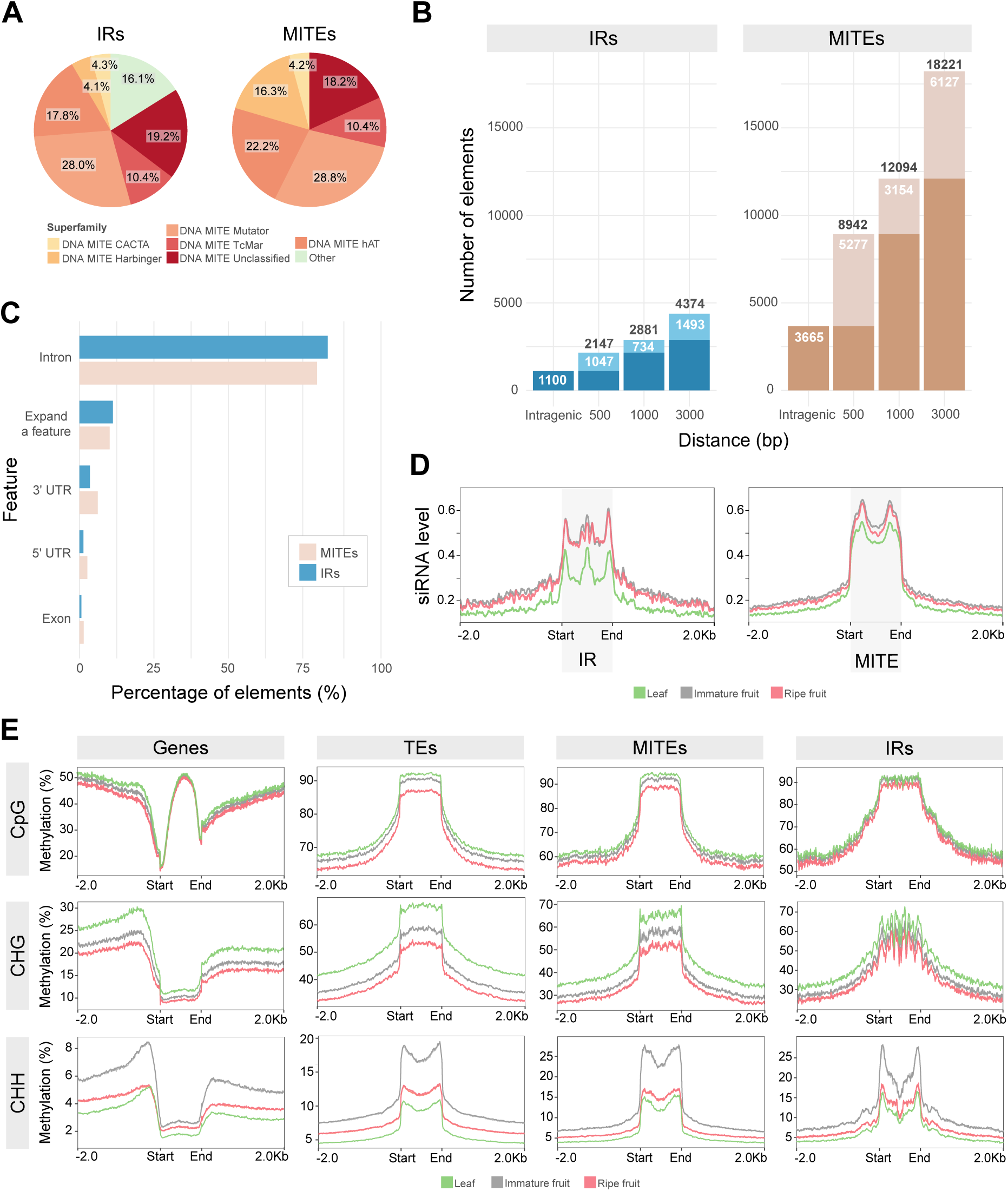
Characterization of TE-Derived IRs and MITEs in *F. vesca*. **(A)** Proportion of the identified IRs (left) and annotated MITEs (right) classified by TE superfamily in *F. vesca.* **(B)** Number of IRs (left) and MITEs (right) detected in the *F. vesca* v4.0.a2 reference genome, counting elements located within gene bodies (‘intragenic’) or within the upstream or downstream regions defined as 0–500 bp (‘500’), 501–1000 bp (‘1000’), and 1001–3000 bp (‘3000’) of annotated genes (the counts for these flanking regions also include any intragenic elements). Black numbers indicate the total counts of IRs and MITEs, while white numbers show the non-cumulative counts. **(C)** Number of IRs and MITEs that are entirely within a single gene feature (‘exon’, ‘intron’, ‘3′-UTR’, or ‘5′-UTR’) or that span more than one gene feature as indicated. **(D)** Metagene profile of 24-nt siRNAs mapping to all IRs and MITEs in leaf (green), and immature (gray) and ripe fruits (pink). Plots show MITEs/IRs scaled from the start to the end plus 2,000 bp on each side. sRNA-seq replicates are plotted together. **(E)** Metagene profile of CpG, CHG, and CHH DNA methylation of all genes, TEs, MITEs, and IRs in leaf (green), immature (shades of gray), and ripe fruits (shades of pink). Plots show Genes, TEs, MITEs, and IRs scaled from the start to the end plus 2,000 bp on each side. Individual BS-seq replicates are plotted.

Most of these identified IRs (4,374, ∼80%) were found near or within annotated protein-coding regions (Figure 3B). Of these, 1,047 IRs are situated 500 bp from a gene, 734 between 501 bp and 1,000 bp, and 1,493 between 1,001 bp and 3,000 bp (Figure 3B). In total, 14% of *F. vesca* genes harbor an IR within these ranges. Similarly, of the 26,060 MITEs annotated (including fragmented elements), 18,221 MITEs were located near or within annotated protein-coding regions. Specifically, 5,277 MITEs were located within 500 bp of a gene, 3,154 between 501 bp and 1,000 bp, and 6,127 between 1,001 and 3,000 bp (Figure 3A). In total, 51% of *F. vesca* genes harbor at least one MITE or IR within 3,000 bp of their coding regions. Notably, 1,100 IRs (∼20%) and 3,665 MITEs (∼14%) are fully located within protein-coding genes, primarily in introns (Figure 3B-C), representing approximately 8% of all coding genes in the *F. vesca* genome. TE insertions in introns can have a significant regulatory effect. For example, in maize, insertion of a CACTA TE derived element into the *Par1* gene creates an epiallele whose methylation status directly affects gene expression and plant architecture [69].

The detected IRs ranged in size from 100 bp to 700 bp (the length limit of our analysis), with an average length of 445 bp. The annotated MITEs exhibited a broader size distribution, ranging from 51 bp to 2,120 bp, with an average length of 261 bp. An overrepresentation of elements between 100 bp and 500 bp (73%) was observed (Figure 1D), values aligning with the typical size range of MITEs in plants [70].

### Most MITEs and IRs in *F. vesca* are epigenetically active

Upon transcription, MITEs and IRs can be processed into siRNA that trigger non-canonical RdDM, leading to the regulation of neighboring genes and turning these TEs into autonomous regulatory elements. To identify potentially active regulatory MITEs and IRs, we used sRNA and bisulfite (BS) sequencing to determine whether the identified MITEs and IRs produced 24-nt siRNAs and triggered DNA methylation—clear indicators of active and potentially regulatory TEs. Three distinct tissues and stages were examined: leaf, and fruit receptacle at both white (immature) and red (ripe) developmental stages (Figure 3D, Figure S5).

Our analysis revealed that 66% of MITEs and 77% of IRs produced siRNAs in at least one of the tested tissues (Table S2). Notably, approximately 80% of these elements are located within 3 kb of a gene, which is consistent with previous reports showing that gene-hosted MITEs and IRs are commonly cotranscribed by RNA polymerase II (Pol II) and are epigenetically active [64]. Metagene analysis of 24-nt siRNAs mapping to the MITEs and IRs showed a characteristic two-peak profile, indicating that the siRNAs are produced from the terminal inverted repeats (TIR) through DLC3 processing (Figure 3D). This behavior was particularly pronounced for IRs located within 1,000 bp of genes. In contrast, for MITEs, this pattern persisted even for elements located farther away from genes, although their expression levels decreased with increasing distance (Figure S5A).

BS-sequencing metagene analysis revealed that annotated TEs, MITEs, and IRs are methylated under all tested conditions (Figure 3E, Figure S5B). Gene-associated MITEs and IRs, particularly those closest to genes, showed a signature two-peak CHH methylation pattern, consistent with the siRNA production profile and RdDM-dependent methylation. Moreover, siRNA levels and CHH methylation decreased as the distance between MITEs or IRs and the associated gene increased (Figure S5B). This distance-dependent epigenetic activity of MITEs and IRs is consistent with previous reports showing greater transcriptional and regulatory activity of elements closer to a gene [64]. This observation supports the idea that 24-nt siRNAs derived from MITEs and IRs near protein-coding genes initiate a non-canonical RdDM pathway likely dependent on Pol II transcription.

In contrast, methylation in the CpG and CHG contexts remained relatively stable regardless of the distance between TEs, MITEs, or IRs and nearby genes (Figure S5B). When these elements were located farther from genes, the surrounding genomic context showed higher CpG and CHG methylation levels, likely reflecting the heterochromatic nature of these regions. Furthermore, CpG and CHG methylation levels were higher in leaves compared to ripe fruits, contrasting with the siRNA production profile and consistent with their independence from siRNA activity (Figure 3D-E, Figure S5B). This contrasts with immature and ripe fruits, which showed higher CHH methylation, suggesting that RdDM activity is required for dynamic regulation of TEs during fruit development and maturation. These differences underscore the distinct epigenetic regulation of transposons across tissues, with stable maintenance of CpG and CHG methylation in somatic tissues and active RdDM-driven CHH methylation in reproductive tissues. This tissue-specific regulatory pattern also reflects the dynamic role of TEs in gene regulation and genome plasticity during fruit ripening and development.

### MITEs and IRs exhibit context-dependent regulatory signatures at the genome-wide level

To investigate the potential regulatory roles of gene-hosted MITEs and IRs in gene expression, we performed a differential accumulation analysis of 24-nt siRNAs derived from these elements as a proxy for MITE differential expression and processing. We compared siRNA profiles between leaves and fruits at two ripening stages (immature and ripe) in *F.vesca*. The analysis revealed a distinct production of 24-nt siRNAs derived from MITEs and IRs across these conditions, considering thresholds of |log2FC| ≥1 and adjusted p-value ≤0.05 (Figure 4A). For MITEs, we identified 17 upregulated and 12 downregulated elements in ripe fruits compared to immature fruits, 318 upregulated and 89 downregulated in ripe fruits compared to leaves, and 89 upregulated and 349 downregulated in leaves compared to immature fruits. Similarly, for IRs, 4 elements were upregulated and 3 downregulated in ripe fruits compared to immature fruits, 144 upregulated and 28 downregulated in ripe fruits compared to leaves, and 26 upregulated and 119 downregulated in leaves compared to immature fruits. This approach allowed us to identify MITEs and IRs actively producing siRNAs in a specific sample, which may serve as potential regulators of gene expression in a tissue and stage-specific manner.

**Figure 4.**
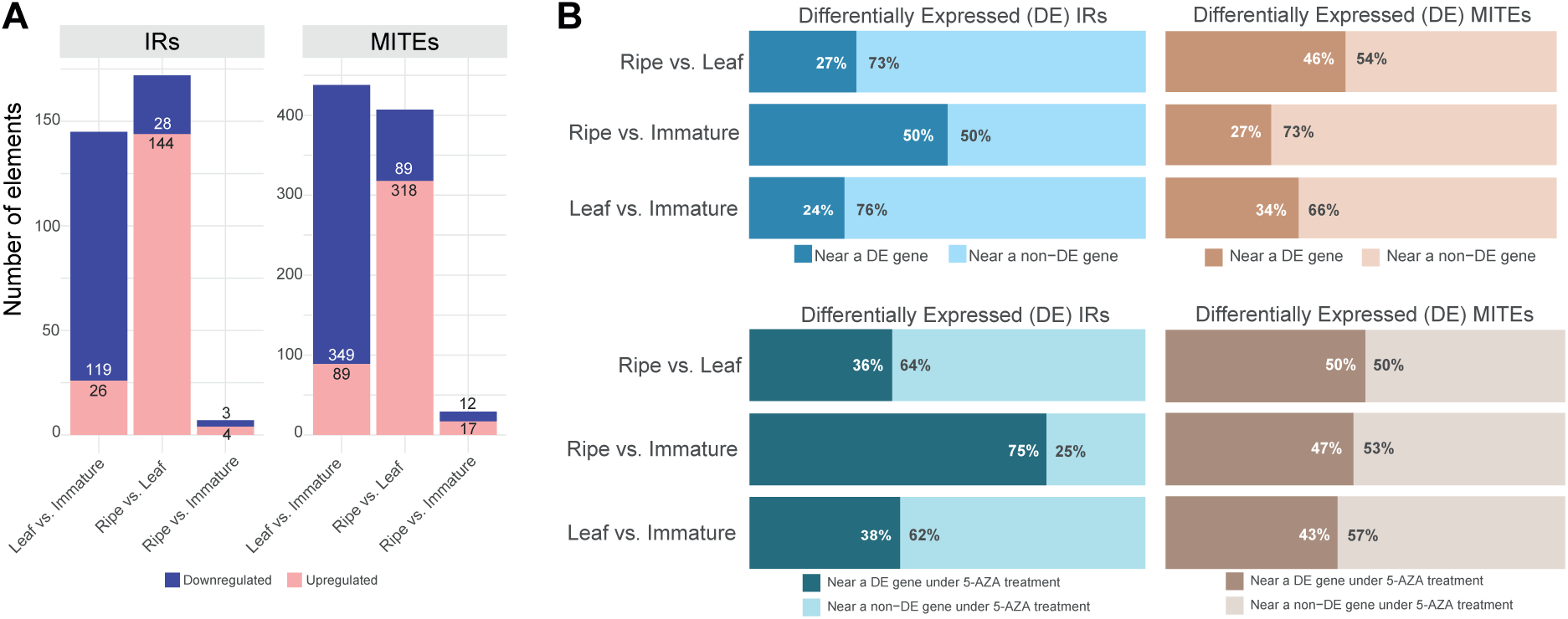
Regulatory role of MITEs and IRs in gene expression. **(A)** Differential accumulation analysis of 24-nt siRNAs derived from IRs and MITEs across leaf tissue, immature and ripe fruits, using thresholds of |log2-fold change| ≥ 1 and an adjusted p-value (padj) ≤ 0.05. **(B)** Differential accumulation analysis of 24-nt siRNAs derived from IRs (left) and MITEs (right) elements across leaf tissue, immature and ripe fruits, using thresholds of log2-fold change (FC) ≥0.5 and adjusted p-value (padj) ≤0.05. The percentages of elements located within 3,000 bp of either differentially expressed genes (DE) or non-DE genes are shown.

We then identified genes located within 3,000 bp of the differentially expressed (DE) MITEs and IRs and evaluated whether these genes also exhibited differential expression under the same conditions (Table S3, Table S4). More than 24% of DE MITEs or IRs were located near DE genes (Figure 4B), with this proportion increasing to 50% during fruit ripening. We also found a, a comparable proportion of non-differentially expressed (non-DE) MITEs and IRs located near DE genes across all conditions analyzed. This indicate that the MITE/IR specific regulation is not necessarily coupled to the transcriptional behavior of the neighboring gene. These cases may parallel our previous observations at the *HaWRKY6* locus in sunflower, where tissue-specific gene regulation is mediated by a MITE-triggered chromatin loop, but the regulatory outcome is controlled by differential DNA methylation at a second anchoring region rather than at the MITE itself, whose expression remains largely stable across tissues but is a *sine-qua-non* requirement for the loop formation [66].

To further assess whether genes associated with MITEs/IRs display distinct transcriptional behavior, we compared the probability of differential expression among genes located near MITEs/IRs, genes associated with non-MITE TEs, and genes in TE-free regions. Genes located in the vicinity (3 kb) of non-MITE TEs showed a significantly lower probability of being differentially expressed compared to genes in TE-free regions (Fisher’s exact test, OR = 0.74, p = 3.7 × 10^−15^), which is expected given that most TE classes are typically located in repressive genomic contexts characterized by reduced transcriptional activity. In contrast, MITE/IR-proximal genes displayed a significantly higher likelihood of differential expression (OR = 1.13, p = 3.3 × 10^−5^), indicating that MITE/IR elements are associated with a distinct regulatory landscape. One limitation in this analysis is that the MITE effect caused by triggering short-range chromatin loops not necessarily work as an on/off switch but affect transcriptional dynamics, transcript structure, or regulatory responsiveness without necessarily producing large expression fold-changes detectable at the genome-wide level [67].

MITEs and IRs have been also reported to act as enhancers or silencers of gene expression by generating cis-regulatory elements [71, 72] or, by modulating local chromatin organization, altering the gene product without changing overall expression levels (e.g., by triggering alternative splicing or termination) [67]. Their regulatory impact depends on multiple factors, including their genomic position relative to the gene, their interaction with regulatory elements such as transcription factors or epigenetic readers, and the way they reshape chromatin architecture, all of which can vary between conditions.

Given the established role of 24-nt siRNAs in mediating RdDM and chromatin remodeling, we next integrated siRNA accumulation with DNA methylation and gene expression data to redefine the identification of MITE/IR-gene pairs with regulatory potential. As a proof-of-concept, we focused on fruit ripening and analyzed publicly available mRNA-seq data from strawberries treated with the DNA methylation inhibitor 5-azacytidine (5-AZA) (Figure 4B, Table S5). Over 35% of DE MITEs/IRs were located near genes that also exhibited significant expression changes upon 5-AZA treatment, with this proportion exceeding 50% during the ripening stage. While this overlap alone does not demonstrate causality, the concordant response of siRNA production, DNA methylation perturbation, and gene expression changes supports the prioritization of specific loci for downstream analysis.

### A MITE Mutator element modulates FvAAT1 expression through epigenetically driven chromatin reorganization

Based on above criteria, we examined individual loci in detail as representative case studies. We focused on elements located near genes with established roles in fruit ripening and associated metabolic or developmental processes, which are highlighted in bold in Supplementary Tables S3 and S4. These case studies are intended to illustrate plausible regulatory configurations supported by epigenetic and transcriptional signals, and are not intended to imply global enrichment or uniform behavior across all analyzed loci.

During fruit ripening we found 29 up- or downregulated MITEs and 7 up- or downregulated IRs near genes associated with fruit ripening, metabolic or developmental processes (Figure 4A, Figure 5A). Among them, we identified a 416 bp element which corresponds to a MITE Mutator, that showed a 12-fold higher siRNA production at the ripening stage (Table S3). This MITE is located 1,915 bp upstream of the gene *FvAAT1* (FvH4_7g18570) (Figure S6), which encodes a putative ortholog to alcohol acyl transferase SAAT, a key enzyme responsible for the biosynthesis of aroma volatile esters in *F. × ananassa [45]*. Consistent with its functional role, *FvAAT1* displayed substantial upregulation during fruit ripening; at the ripe stage, gene expression was more than 1,700 times higher than at the immature stage (Table S5), aligning with previous findings that underscore its importance in producing characteristic ester-based aroma compounds during ripening [73, 74]. This gene is also upregulated during the turning stage in *F. × ananassa* compared to *F. vesca*, emphasizing its importance in commercial fruit quality traits [75]. By contrast, treatment with 5-AZA resulted in a fivefold reduction in *FvAAT1* expression (Table S5), suggesting that the identified epigenetically active MITE can influence its expression.

**Figure 5.**
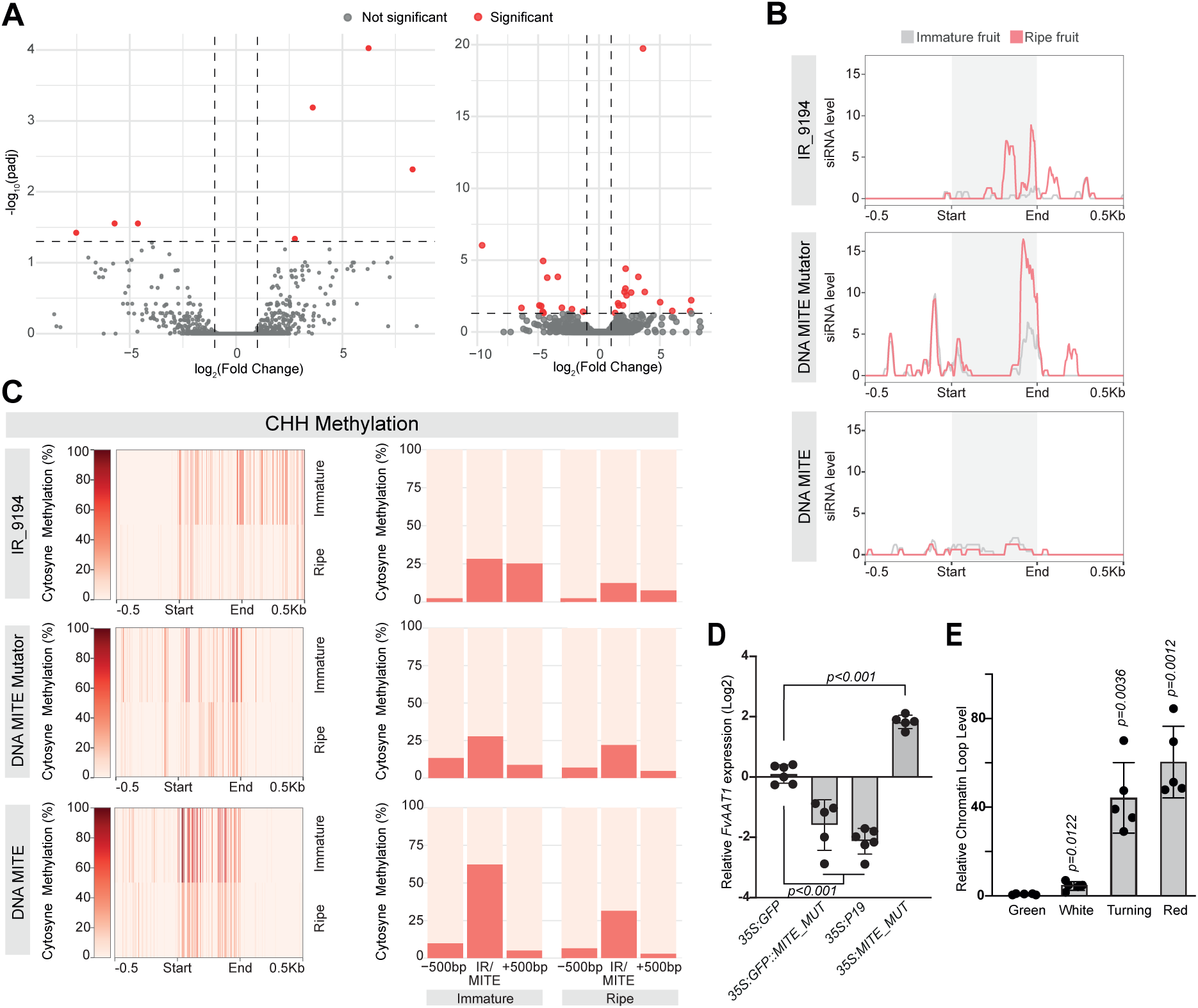
A *FvAAT1*-hosted MITE control the gene expression through epigenetically driven chromatin reorganization. **(A)** Volcano plots showing the log2FC of the differential expression analysis for IRs (left) and MITEs (right) between immature and ripe stages. Red points represent elements with an adjusted p-value ≤0.05. **(B)** 24-nt siRNAs mapping to IR_9194 and the two MITEs near FvH4_7g18550 and FvH4_7g18570 genes (highlighted by a gray box) in immature (gray) and ripe (pink) fruits. Each element is scaled from start to end plus 500 bp on each side, and all sRNA-seq replicates are plotted together. **(C)** Cytosine methylation in the CHH context for *IR_9194*, MITE Mutator, and MITE, including 500-bp flanking regions, in immature and ripe fruits. The average of individual BS-seq replicates is plotted for each condition. Average percentage of CHH methylation in each element and its 500-bp flanking regions. **(D)** *FvAAT1* expression in transiently transformed strawberry receptacles. RNA levels are expressed relative to the control samples (35S:GFP) plants. Values of n = 6 biologically independent samples. Data are presented as the mean values ± s.d. P values were calculated with a two-tailed unpaired t-test with Welch’s correction. **(E)** Chromatin conformation capture (3C) analysis showing the interaction frequency of MITE Mutator with the FvAAT1 locus in immature and ripe fruits as measured by 3C–qPCR. Values of n = 4 biologically independent samples are expressed relative to green immature stage. Data are presented as the mean values ± s.d. P values were calculated with a two-tailed unpaired t-test with Welch’s correction.

In the same genomic region, we identified another MITE located 1,064 bp downstream of *FvAAT1* and an IR element (IR_9194) located 4,761 bp upstream of *FvAAT1* and 3 bp downstream of FvH4_7g18550, a gene encoding a papain family cysteine protease implicated in protein degradation and defense responses during fruit ripening (Figure S6). This gene showed a marked expression increase (∼105-fold) in ripe fruit compared to immature fruit (Table S5), mirroring patterns observed in other plant species in which cysteine proteases have been proposed to regulate ripening processes [76–78].

We then evaluated siRNA production and CHH methylation patterns around these elements to gain further insights into potential epigenetic regulation (Figure 5B and 5C). We observed that IR_9194 and especially the MITE Mutator produce 24-nt siRNA with higher levels in <u>fruit ripening</u>, consistent with active <u>elements</u> (Figure 5B, top and middle panel, respectively). This increase in siRNA production in mature fruits coincides with a CHH methylation signal (Figure 5C). Interestingly, the MITE located downstream of FvH4_7g18570 displayed the highest methylation levels at its TIRs during immature fruit stage, albeit we did not observe a significant accumulation of siRNA (Figure 5B and 5C, bottom panel). This may indicate that this MITE might be in a permanently silenced state, in contrast to IR_9194 and the MITE Mutator, which remain epigenetically active.

Although such MITE appears to be transcriptionally inactive, the increase in DNA methylation at this element may serve as an “anchor point” in conjunction with the more dynamic MITE Mutator element to trigger chromatin looping and control *FvAAT1*, as observed in other systems [66]. To test the hypothesis that the promotion of a chromatin loop between these two elements impact the expression of FvAAT1, we designed an experiment to transiently modulate the epigenetic status of the MITE Mutator. To do so, we infiltrated *F. vesca* fruits with a construct either overexpressing the viral silencing suppressor P19, which reduces siRNA activity and thus RdDM, or an additional copy of the MITE Mutator under the 35S promoter to increase the production of its specific siRNAs *in trans* and thus induce methylation at this element. We have previously used these strategies to induce or repress MITEs activity [65, 66]. We then measured the expression of *FvAAT1* in infiltrated fruits and mock infiltrated controls. Consistent with the proposed regulatory effect of the MITE Mutator over *FvAAT1,* we observed a correlated expression of the gene when RdDM at the MITE was induced or repressed (Figure 5D). The activation *in trans* of the MITE Mutator correlated with a significant increase in the expression of *FvAAT1* (Figure 5D). Conversely, the epigenetic inactivation of the MITE Mutator, achieved by transiently overexpressing either a MITE Mutator sponge sequence fused to GFP, which sequesters the MITE-derived siRNAs, or the viral silencing suppressor P19, which nonspecifically sequester siRNAs, correlated with a significant repression of the gene *FvAAT1* (Figure 5D). Furthermore, using a chromatin conformation capture (3C) approach, we observed a ripening stage dependent change in chromatin organization, with the MITE Mutator acting most likely as the anchor point (Figure 5E). In this sense, a chromatin loop encompassing the entire loci was increasingly detected with the maturation of the fruits, suggesting a positive effect on gene transcription, as previously reported for example during Pol II reclining happening on gene looping events [67]. Given the apparent regulatory importance of the MITE Mutator, we assessed its conservation in the orthologous genomic regions of the commercial strawberry species *F. chiloensis* and *F. × ananassa* through sequence alignment. In *F. chiloensis*, a copy of the element, situated 1,674 bp upstream of the *FvAAT1* homologue was detected, displaying 89.4% sequence identity (Figure S7A, left). A similar pattern was observed in *F. × ananassa* (Camarosa genome assembly), where a copy of the MITE associated with *FaAAT1* exhibits 95.7% sequence identity (Figure S7A, right). Analysis of the predicted secondary structures using RNAfold for the three elements revealed the formation of a long dsRNA hairpin structure, suggesting that they are likely DCL3 substrates and will function similarly to those observed in *F. vesca* (Figure S7B). These findings underscore the evolutionary conservation of this MITE near key ripening-related genes across different *Fragaria* species, potentially reflecting a conserved regulatory function in fruit quality traits.

We identified additional IR/MITE elements adjacent to genes with key roles in fruit development and metabolism, suggesting potential regulatory contributions through small RNA production, DNA methylation, and gene expression changes (Table S4 and Table S5). For example, a 439 bp IR element (IR_8440) exhibiting approximately a 4.5-fold increase in expression in red fruit compared to leaf tissue (Table S4) and pronounced CHH methylation in fruit was identified 1,972 bp upstream of the SEPALLATA-like MADS-box gene FvH4_6g46420 (Figure S8A). This IR is adjacent to a MITE Harbinger. A MITE hAT located downstream of FvH4_6g46420 displays relatively constant expression across tissues, suggesting it could serve as a potential “anchoring” region for 3D chromatin configurations. The FvH4_6g46420 gene itself is significantly upregulated in white fruit vs. leaf tissue, consistent with prior studies underscoring the role of its ortholog in *F. × ananassa*, *FaMADS9*, which is involved in strawberry fruit development and ripening [79]. Silencing of *FaMADS9* in *F. × ananassa* delays or inhibits normal ripening and alters both auxin and ABA metabolism, implicating this MADS-box transcription factor in pleiotropic regulatory networks [80, 81]. Furthermore, *FaMADS9* is downregulated under AZA treatment, highlighting the broader interplay between epigenetic modifications and fruit ripening pathways [82].

Additional cases reinforce the potential role of IRs and MITEs in regulating nearby gene expression. A MITE hAT (334 bp) is located near FvH4_2g30230, annotated as molybdate transporter 1 (Figure S8B). Molybdenum is an essential trace element in plant nutrition and physiology, and its effects on strawberry flavor and fruit quality are well documented [83]. The upregulation of this gene in ripe fruit and its downregulation in 5-AZA-treated strawberries, along with the distinct expression profiles of the MITE in leaf tissue and in fruits at the white and red ripening stages, together with localized methylation and siRNA mapping, further support a role for IR-associated epigenetic control. Moreover, IR_6046 (448 bp) is located upstream of FvH4_5g23180 (Figure S8C), encoding a probable xyloglucan endotransglucosylase/hydrolase (XTH30), which is critical for cell wall remodeling during fruit softening [84, 85]. Differential expression of both the element and XTH30, alongside distinct siRNA clusters in leaf versus fruit ripening conditions, aligns with a mechanism involving dynamic chromatin organization. Fruit softening is a critical factor affecting the marketability and economic value of fruits; therefore, targeting cell wall-related genes, including XTHs, may offer an effective strategy to delay this process and extend postharvest shelf life.

Taken together, these observations underscore the potential of IR and MITE elements to modulate gene activity through epigenetic pathways, thereby influencing fruit maturation and quality traits. An idea strongly supported by the validations made over the *FvAAT1* loci.

### Discovery of TE Polymorphisms in resequenced *F. vesca* genomes

Due to their inherent mobility, TEs frequently vary between natural populations in copy number and genomic position, producing insertion or deletion polymorphisms [18]. A TE insertion polymorphism (TIP) occurs when a TE is present in an accession but absent from the reference genome, whereas a TE absence polymorphism (TAP) represents a TE found in the reference but missing in other accessions. Because many TEs harbor their own regulatory sequences, they can strongly influence the expression of nearby genes, often more than other types of structural variation [36, 86]. Moreover, our previous results indicate that MITEs and IRs potentially exert a regulatory effect upon insertion into intergenic regions proximal to genes [64]. Historically, genetic diversity analyses have primarily focused on SNPs, while TE insertion/deletion polymorphisms remain an underexplored source of genomic variation [87–90]. TEs activity is often influenced by stress conditions and many causative TE insertions result from recent movements and are not effectively tagged by SNPs, meaning TIPs can be used as direct markers to uncover these adaptive genetic factors [32, 91]. As such, they hold the potential to affect phenotypes and key biotechnological traits in nutritionally and commercially important plant species.

We analyzed 210 resequenced *F. vesca* genomes from a European germplasm collection, representing the species’ continental diversity [44] (PRJNA1018297), to study natural variation in TEs, MITEs, and IRs using TEPID [90] (Figure S9). A total of 8,184 TE, MITEs and IRs insertions (TIPs) and 11,345 deletions (TAPs) were identified across the population (Table 2). As reported for other plant species [32, 92, 93], the TE landscape was prevalent in rare variants; 84% of TE insertions and 88% of IR insertions exhibited a MAF below 5%. This strong skew toward rare alleles suggests that these insertions are either evolutionarily recent or are being actively purged by purifying selection to maintain genomic integrity. Notably, deletion polymorphisms reached higher frequencies in the population than insertions; for instance, 28.7% of TE deletions were classified as common (MAF ≥ 5%) compared to only 15.8% of insertions. This disparity suggests that many deletions represent the loss of ancestral elements present in the reference genome that have not reached fixation in the European germplasm. Among the identified TE insertions, 35% belonged to Class I and 65% to Class II, closely matching the general TE distribution observed in the reference genome (33% and 67%, respectively). A similar trend was found for deletion polymorphisms, where Class I elements accounted for 35% and Class II for 65% of the total events Insertion polymorphisms were particularly enriched for LTR/Gypsy, hAT, Mutatorand MITEs elements, suggesting recent bursts of activity that have not yet spread across the population [94] (Figure 6A). Conversely, LTR/Gypsy deletions were notably frequent among common variants, indicating a significant turnover of these retrotransposons in specific lineages.

**Figure 6.**
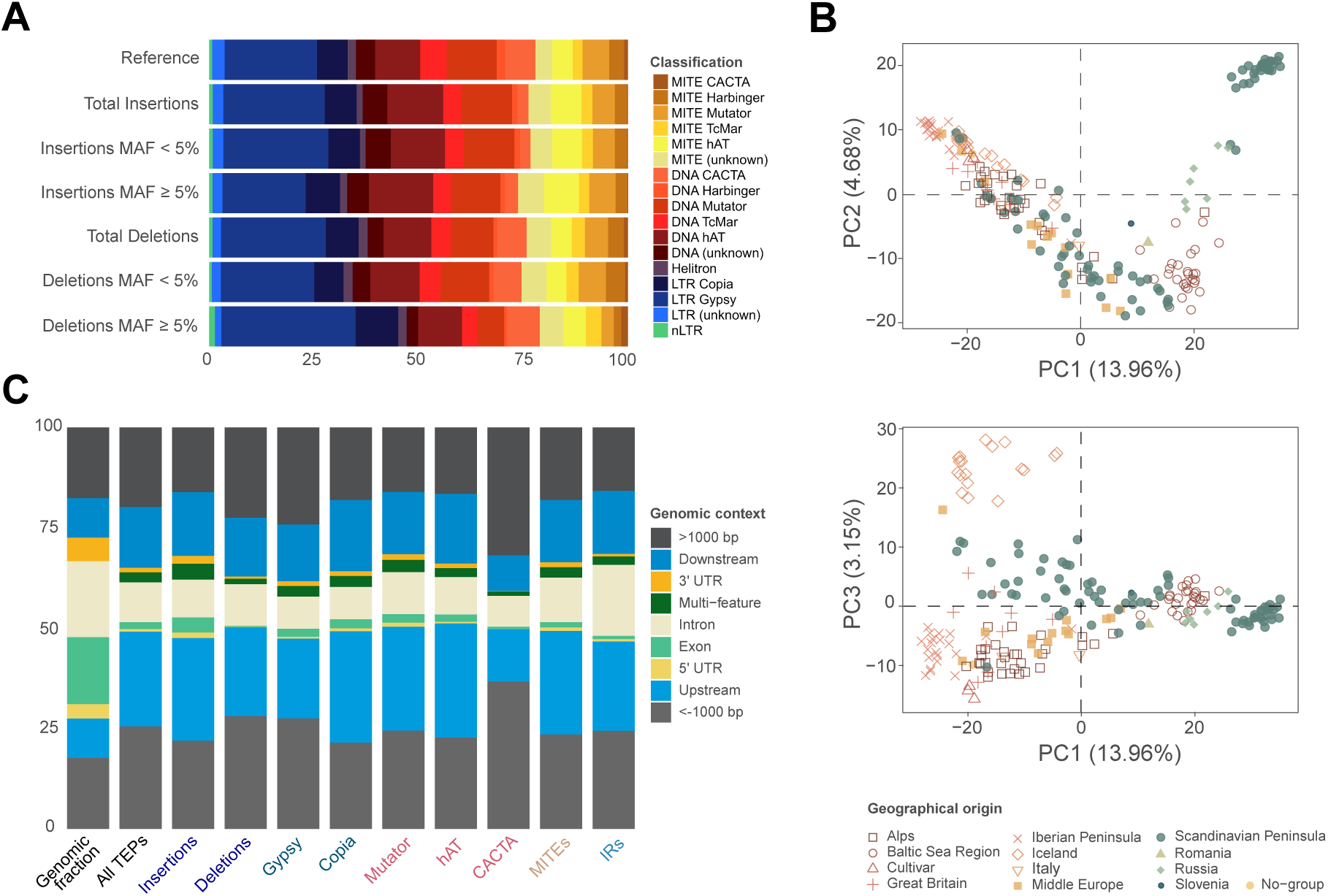
Distribution of insertion and deletion polymorphisms in the *F. vesca* v4.0.a2 annotation. (A) Proportion (%) of identified polymorphisms by TE superfamily. Rare TE polymorphisms are indicated by minor allele frequency (MAF) < 5% and frequent TE polymorphisms by MAF ≥ 5%. (B) Principal Component Analysis (PCA) based on TE insertion and deletion polymorphisms (MAF ≥ 5%). The geographic origin of the analyzed F. vesca accessions is indicated according to Urrutia et al [95]. **(C)** Distribution of polymorphic elements over genic features, including 5’ UTR, Exons, Introns, and 3’ UTR (UTR: unstranslated transcribed region; Multi-feature: polymorphisms overlpaping multiple annotations). Stacked bar plots represent the relative proportion of Insertions, Deletions, and specific TE superfamilies. Polymorphisms located within 1000 bp of a gene are shown in shades of blue, representing Upstream and Downstream regions. Intergenic polymorphisms located more than 1000 bp away from those regions are indicated in grey (<-1000 bp and >1000 bp). The ‘Genomic fraction’ bar represents the percentage of base pairs occupied by each feature in the reference genome.

**Table 2.** Polymorphisms detected by TEPID for each element annotation. MAF ≥ 5% refers to polymorphisms with a minor allele frequency above 5%. The percentage shown represents the proportion of each value relative to the total number of polymorphic elements detected for each element and each type (insertion/deletion).

| Polymorphisms |  | TEs |  | MITes |  | IRs |  |
| --- | --- | --- | --- | --- | --- | --- | --- |
|  |  | Amount | % | Amount | % | Amount | % |
| Insertions | Total | 5974 | 100.0 | 1859 | 100.0 | 351 | 100.0 |
|  | MAF ≥ 5% | 946 | 15.8 | 336 | 18.1 | 42 | 12.0 |
|  | MAF < 5% | 5028 | 84.2 | 1523 | 81.9 | 309 | 88.0 |
| Deletions | Total | 7855 | 100.0 | 2507 | 100.0 | 983 | 100.0 |
|  | MAF ≥ 5% | 2256 | 28.7 | 601 | 24.0 | 197 | 20.0 |
|  | MAF < 5% | 5599 | 71.3 | 1906 | 76.0 | 786 | 80.0 |

A principal component analysis (PCA) based on the identified TE polymorphisms revealed a clear subpopulation structure (Figure 6B), mirroring the geographical clustering previously reported using SNP-based markers [44]. In the TE-based PCA, the PC1 and 2 captured ∼19% of the variance, similar to the SNP-based PCA by Toivainen et al. (2026) [44]. This provides strong confidence in the accuracy and reliability of our detected polymorphisms, as both genomic features capture the same fundamental evolutionary signals. Our TE-based analysis also reflected the differentiation reported in the *F. vesca* volatilome by Urrutia et al. [95], such as the distinct separation of the Icelandic and Baltic Sea clusters. The TE-based PCA also showed a clear resolution for the Iberian and Scandinavian peninsulas, and a separation of Western and Southern European accessions from Northern and Eastern populations. In Scandinavia, this is particularly evident and may be linked to the admixture of western and eastern post-glacial colonization routes [44]. These results suggest that TE-derived structural variation provides a highly sensitive molecular signature for uncovering fine-scale population structure in *F. vesca*.

The genome-wide distribution of insertion and deletion polymorphisms in *F. vesca* revealed a clear non-random pattern shaped by genome architecture and selective constraints (Figure 6C). Insertions and deletions exhibited partially distinct patterns. Deletions showed a stronger avoidance of exonic regions than insertions, in line with their greater potential to compromise gene integrity (Figure S10A). Across all TEs superfamilies, polymorphisms were strongly depleted within gene bodies, with the most pronounced avoidance observed in exons and untranslated regions (UTRs), consistent with strong purifying selection acting against disruptive structural variation in functional elements (Figure 6C, Figure S10B). Introns showed a comparatively milder depletion, indicating a higher tolerance for structural variation relative to coding and regulatory regions. In contrast, polymorphisms were consistently enriched in gene-flanking regions, where chromatin is typically less condensed, likely facilitating access for the TE mobilization machinery (Figure 6C, Figure S10). Similar patterns have been observed in other species, where TE insertions tend to occur in gene-rich regions, although variations can occur at both the TE family and species levels [36, 94, 96]. Overall, more than half of all identified polymorphisms were located within genes or their immediate flanking sequences (up to 1,000 bp upstream/downstream). Specifically, MITEs and IRs showed a preferential accumulation within promoter regions (Figure 6C, Figure S10B). This high frequency in upstream regions and to some extent downstream, suggests a significant potential for these elements to influence gene expression, as observed in the results described above or in other crop species, like *Brassica rapa* and *Solanum lycopersicum* [36, 37].

Taken together, these results suggest that the landscape of TE polymorphisms in *F. vesca* is characterized by a high turnover of elements within regulatory proximity to genes. The robust population structure resolved by these variants underscores the role of TEs as dynamic footprints of post-glacial colonization. The high frequency of rare variants suggests that most transposable element polymorphisms (TEPs) sare subjected to purifying selection or occurred recently in certain lineages. Moreover, these TEPs occur preferentially in promoter regions. Collectively, this represents a significant and underexplored reservoir of genetic diversity with the potential to impact complex phenotypic traits in European woodland strawberry germplasm.

### Linking TE polymorphisms to phenotypic traits in woodland strawberry fruits

Several phenotypic traits, including fruit dimensions (length, width, volume) and the concentrations of 99 volatile secondary metabolites (the volatilome), were previously measured in a collection of 116 and 154 wild *F. vesca* accessions across two independent harvest seasons (H16 and H17) [95]. Since the genomes of these accessions were included in our polymorphism analysis, we leveraged the detected TEPs to perform a genome-wide association study (GWAS) to investigate the impact of TE and IR natural diversity in those strawberry fruit traits.

Several phenotypic traits, including fruit dimensions (length, width, volume) and the concentrations of 99 volatile secondary metabolites (the volatilome), were previously measured in a collection of 116 and 154 wild *F. vesca* accessions across two independent harvest seasons (H16 and H17) [95]. Since the genomes of these accessions were included in our polymorphism analysis, we leveraged the detected TEPs to perform a genome-wide association study (GWAS) to investigate the impact of TE and IR natural diversity in those strawberry fruit traits.

For the volatilome dataset, 68 distinct TEPs were significantly associated (5% Bonferroni threshold) with 40 VOCs across harvests H16 and H17 resulting in 78 significant trait–polymorphism associations (Table S7). None of these associations corresponded to TEP directly impacting the coding sequence of genes known to be involved in strawberry VOC metabolism; this is consistent with our observation that only a very small fraction of these variants occur in exonic regions and tend to occur more frequently in intergenic regions proximal to genes where they could exert a regulatory effect (Figure 6C). This is also suggested in the results in this work presented, in which we observed the regulatory effect of different MITEs on the expression of *FvAAT1*.

Considering our results, we next focused on statistically significant associations that exhibit potential biological relevance regulating the genes in their vicinity based on current knowledge, specially focusing on MITE or IR polymorphisms. For instance, a MITE Mutator insertion was associated with 6-methyl-5-hepten-2-one (sulcatone) on chromosome Fvb3 (*P* = 5.6512 × 10^−6^; PVE = 9.20%) (Figure 7A-B, Figure S12A). Sulcatone is a common apocarotenoid that originates from lycopene cleavage [97]. Apocarotenoids contribute to the aromas of different flowers and fruits and are highly valued by the flavor and fragrance industry [97]. Unsaturated methyl ketones, such as sulcatone, have been proposed to act as highly reactive VOCs with defensive and pesticidal properties and have been associated with unpleasant taste in fruits affected by powdery mildew disease [98–100]. Accessions carrying this insertion (n = 34) showed lower 6-methyl-5-hepten-2-one levels in harvest H17 (Figure 7C). Interestingly, the insertion is located in a gene-dense genomic region (FvH4_3g02700, FvH4_3g02710, FvH4_3g02720, FvH4_3g02730 and FvH4_3g02731 genes) enriched in stress- and ROS-related signalling components and redox enzymes, including LEA/NDR1-like (LEA-2) proteins, NDR1 a secretory class III peroxidase, a cysteine-rich receptor-like kinase (CRK-like), and an LpxD-like N-acyltransferase. Class III peroxidases act as major apoplastic regulators of H₂O₂ homeostasis and oxidative reactions, shaping the local redox environment that feeds into downstream oxidative volatile formation [101]. In parallel, NDR1 is a central component of plant defence signalling, and CRK family kinases are widely implicated in ROS-associated signal transduction during stress responses. The presence of a MITE element in this gene module could impact the regulation of these genes, ultimately affecting the response to oxidative stress and indirectly influencing the oxidative flux required for carotenoid degradation and the sulcanone levels.

**Figure 7.**
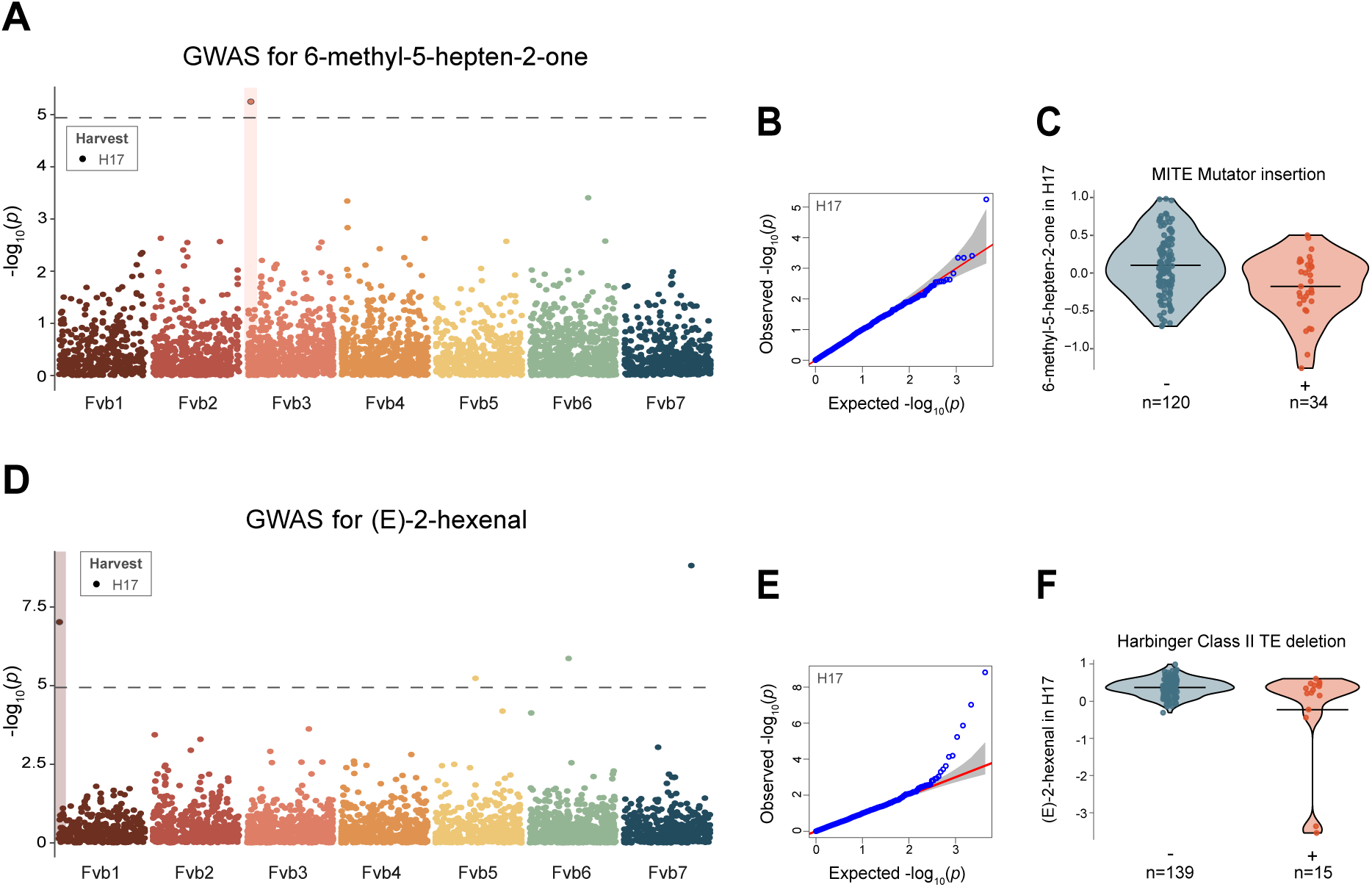
TE polymorphisms linked to volatile compound variation in a collection of *F. vesca* accessions. **(A)** Manhattan plot of TEP-GWAS for 6-methyl-5-hepten-2-one in harvest season H17. The dotted gray line marks the significance threshold determined by the Bonferroni correction, and the significantly associated polymorphism is highlighted. **(B)** Observed and expected distribution of *p* values for TEP-GWAS for 6-methyl-5-hepten-2-one. **(C)** Violin plot showing the relative abundance of 6-methyl-5-hepten-2-one in accessions carrying the MITE insertion (+, coral) versus non-carriers (-, steel blue). Horizontal lines in (-) and (+) indicate the mean. **(D)** Manhattan plot of TEP-GWAS for (E)-2-hexenal in harvest season H17. The dotted gray line marks the significance threshold determined by the Bonferroni correction, and the significantly associated polymorphism is highlighted. **(E)** Observed and expected distribution of *p* values for TEP-GWAS for (E)-2-hexenal. **(F)** Violin plot showing the relative abundance of (E)-2-hexenal in accessions carrying the MITE insertion (+, coral) versus non-carriers (-, steel blue). Horizontal lines in (-) and (+) indicate the mean.

A Harbinger Class II TE deletion was associated with lower levels of the 6-C aldehyde green leaf volatile (GLV) (E)-2-hexenal in H17 (*P* = 9.51 × 10^−8^; PVE = 2%) (Figure 7D-F). GLV are VOCs emitted in response to wounding or plant damage and are responsible for the characteristic “green” aroma of plants (reviewed in [102]). The deletion is located on chromosome Fvb1, 13 kb upstream of a gene annotated as 2-ALKENAL REDUCTASE (FvH4_1g00330). Alkenal reductases facilitate the reduction of double-bonds in aldehydes, including short-chain aldehydes such as (E)-2-hexenal [103, 104]. This gene, or any of its eight downstream paralogs, represents a potential candidate for the conversion of (E)-2-hexenal to hexanal in strawberry. Notably, this locus has not been identified in previous QTL analysis in *F. vesca* or cultivated strawberry [105, 106], highlighting the potential of TEPs-GWAS to identify novel QTLs.

Other associations further suggest linkage between the TEP and the underlying candidate gene. A primary example is methyl 2-aminobenzoate (methyl anthranilate), a precursor of tryptophan involved in primary metabolism and associated with a grape-like aroma in fruits [107]. We identified a significant association with the deletion of a MITE Mutator on chromosome Fvb5 (*P* = 1 × 10^−14^; PEV = 0) (Figure S13A-C). Interestingly, although this variant appears to explain only a marginal amount of phenotypic variance, this locus has been consistently associated with methyl anthranilate levels in octoploid strawberry [108]. Barbey et al. described a QTL at the beginning of chr5-4, the same region where the TEP is located. This site is 380 kb from an L-tryptophan—pyruvate aminotransferase (TAA1) gene (FvH4_5g05880), which utilizes L-tryptophan for auxin biosynthesis; this gene may be related to varying levels of the intermediate methyl anthranilate.

Our analysis also identified a MITE Harbinger deletion on chromosome Fvb3 that was associated with γ-decalactone levels across both harvest’s seasons (*P* = 4.2911 × 10^−6^ in H16 and *P* = 1.4523 × 10^−6^ in H17) (Figure S13D-F, Figure S12B). This variant appears to explain only a marginal amount of phenotypic variance in both harvests. Accessions carrying this deletion (n = 87 and 115 in H16 and H17, respectively number) showed higher levels of γ-decalactone, a key compound previously identified as one was one of the most variable in the *F. vesca* volatilome analysis by Urrutia et al. [95]. In *F. × ananassa*, γ-decalactone is the most abundant lactone [79], reaching maximum levels in fully ripe fruits [79] and is positively correlated with consumer perception of favorable organoleptic traits [95, 109]. The deletion falls within an intron of the gene FvH4_3g30910, annotated as a replication factor-A C-terminal domain protein, which shows no evident functional connection to γ-decalactone. Notably, however, the deletion lies ∼5 kb downstream of FvH4_3g30901, a gene encoding a serine carboxypeptidase-like (SCPL) enzyme. SCPL enzymes belong to the α/β-hydrolase superfamily, functioning as acyltransferases in plant secondary metabolism which include esterases and acyltransferases that act on diverse lipid substrates [110, 111]. Given the functional plasticity of these enzymes, it is likely that many SCPL members involved in fruit specialized metabolism are yet to be characterized. In this context, structural variation affecting chromatin state or *cis*-regulation near the SCPL gene could indirectly contribute to quantitative differences in γ-decalactone.

Taken together, these results suggest that natural polymorphisms in transposable elements, particularly MITEs and IRs, are linked to variation in strawberry fruit volatilome composition, primarily through effects in gene-proximal regions. While these associations do not imply direct causality, they reveal candidate regulatory loci whose effects are shaped by local genomic and chromatin context.

## DISCUSSION

Our study provides a comprehensive analysis of TEs in *Fragaria vesca*, highlighting their role in genome composition, organization, and potential functional impact. By systematically annotating TEs and analyzing their polymorphisms across 210 *F. vesca* accessions, we have gained valuable insights into their evolutionary significance and their potential contributions to gene regulation and phenotypic diversity.

Our TE annotation revealed that TEs occupy ∼30% of the *F. vesca* genome, a proportion consistent with expectations for plant species of similar genome size and slightly higher than the 22% reported in the initial 2011 *F. vesca* genome assembly [41]. Unlike most plant genomes, where LTR retrotransposons are dominant [54] *F. vesca* exhibits a comparable proportion of Class II DNA transposons, with MITEs representing a substantial fraction of intact elements. This enrichment is likely explained by their compact structure and lack of coding regions, which reduce their susceptibility to fragmentation over evolutionary time. In contrast, LTR retrotransposons, particularly the larger elements, appear more frequently fragmented, consistent with higher rates of structural decay. The genome-wide distribution of TEs further reveals distinct patterns among classes: LTRs are enriched in gene-poor, pericentromeric regions, although their relatively low abundance in *F. vesca* may reflect incomplete assembly of centromeric regions, whereas MITEs, IRs, and other DNA transposons are concentrated near or within coding sequences, suggesting a potential role in the regulation of adjacent genes. These observations underscore the unique composition of the *F. vesca* genome and provide a foundation for exploring how TE-mediated regulatory variation contributes to phenotypic diversity in strawberry. One of the most significant findings of our study is the widespread production of small RNAs (siRNAs) from MITEs and IRs in *F. vesca*, supporting their potential regulatory role in gene expression. On average, 71% of these non-autonomous elements produced 24-nt siRNAs in at least one tissue type studied, with a characteristic two-peak siRNA profile indicative of processing through the RdDM pathway. The correlation observed between siRNA production and CHH methylation, particularly in reproductive tissues, underscores the functional relevance of these elements in tissue-specific gene regulation. In contrast, CpG and CHG methylation patterns remained relatively stable, supporting the role of siRNA-driven CHH methylation as a key mechanism controlling TE activity and gene expression in a dynamic and reversible manner. These findings highlight the functional significance of MITEs and IRs as autonomous regulatory elements that modulate gene expression through small RNA-mediated epigenetic modifications. These observations position MITEs and IRs not merely as silencing targets, but as active components of regulatory networks capable of modulating local chromatin states.

While genome-wide analyses revealed that differentially expressed MITEs and IRs are not significantly more likely than non-DE elements to be located near differentially expressed genes, this finding underscores an important conceptual point: TE proximity alone is insufficient to infer regulatory function. This is not entirely surprising as we previously found that one of the most important features of MITE affecting gene expression is not their sequence but their capacity to trigger chromatin loops, which inevitably require an additional anchor point. In a previous study we showed that, if a MITE is expressed, its regulatory function can be modulated by alterations (methylation, accessibility, etc.) on the second anchor point [66]. The lack of global enrichment of differentially expressed genes near DE MITEs or IRs during ripening further indicates that these elements do not act alone. Instead, their regulatory potential appears to emerge from the convergence of multiple features, including tissue-specific siRNA production, local methylation dynamics, transcriptional responsiveness, and the genomic neighborhood.

This concept is exemplified by the detailed analysis of selected loci associated with fruit ripening. At the *FvAAT1* locus, a MITE Mutator element displays coordinated increases in siRNA accumulation and a clear CHH methylation signature during ripening, coinciding with dramatic upregulation of the adjacent gene encoding an alcohol acyltransferase, a key enzyme responsible for volatile ester biosynthesis in strawberry [45, 112, 113]. The upregulation of *FvAAT1* during ripening, coupled with siRNA production from the adjacent MITE, supports a model in which this TE influences gene expression through methylation-dependent chromatin remodeling. Further validation using 5-AZA demethylation assays confirmed that *FaAAT1* expression is methylation-dependent, highlighting the direct regulatory effect of MITEs in *F. vesca* fruit ripening. The conservation of this particular MITE in *F. chiloensis* and *F. × ananassa*, with high sequence identity and structural similarities, suggests that it plays a conserved role in fruit aroma regulation across strawberry species. This discovery adds to the growing evidence that TEs contribute to the evolution of crop traits by introducing regulatory variation that affects key metabolic pathways [7, 36, 114, 115]. It also positions TEs, especially gene-associated MITEs and short IRs, as promising targets for genome editing. The CRISPR/Cas9-mediated removal of a MITE could significantly alter the expression of adjacent genes and the associated phenotypic traits. Genome editing of MITEs also offers the advantage that these elements are non-coding and naturally variable in populations, potentially simplifying regulatory approval.

In addition to their regulatory potential, TE polymorphisms provide an important source of genetic diversity that can influence trait variation [37]. Our GWAS analyses link TE polymorphisms to volatile composition, demonstrating the phenotypic relevance of mobile elements in *F. vesca*. Notably, some of the identified TEPs correspond to previously unreported loci affecting key flavor compounds, such as sulcatone, γ-decalactone, and methyl anthranilate, highlighting the potential of TEP-focused GWAS to uncover novel regulatory loci that are missed by traditional SNP-based approaches. While these associations do not establish direct causality, they provide candidate regulatory elements for functional validation and suggest that integrating TE polymorphisms into breeding programs could expand the spectrum of allelic variation available for trait improvement. Our findings demonstrate that TE-driven variation significantly contributes to the natural diversity of fruit metabolic profiles, with potential implications for breeding strategies aimed at improving fruit aroma and flavor. Furthermore, understanding how TEs modulate gene expression and metabolite production at the molecular level provides valuable insights into the complex regulatory networks underlying crop traits. The high degree of TE polymorphism observed in *F. vesca*, combined with the functional relevance of associated traits, suggests that TEs represent an untapped source of genetic variation for crop improvement.

## CONCLUSIONS

Our findings demonstrate that, as in other plants, TEs in strawberries are not merely passive genomic elements but rather key contributors to genome function, gene regulation, and phenotypic variation. By integrating TE annotation, small RNA profiling, methylation data, gene expression analyses, and TEP-based GWAS, we identify specific TE–gene configurations with plausible regulatory roles in fruit development and quality traits. Although TE proximity alone does not predict regulatory function at the genome-wide level, selected loci illustrate how epigenetically active MITEs and IRs can modulate gene expression in biologically relevant contexts, such as fruit ripening.

This identification of TE-derived regulatory elements and polymorphisms associated with fruit traits provides a valuable resource for future breeding efforts aimed at improving strawberry quality. In particular, the regulatory roles of MITEs and IRs in fruit ripening suggest that targeted epigenetic modifications could be leveraged to optimize fruit quality. The use of genome-editing tools such as CRISPR/Cas9 to precisely manipulate TE-derived regulatory regions may enable the fine-tuning of gene expression without introducing foreign DNA, making it a promising strategy for crop improvement.

Additionally, the conservation of key regulatory MITEs in commercial strawberry species suggests that similar TE-driven mechanisms may underlie important agronomic traits in cultivated varieties. Further comparative genomic studies across *Fragaria* species could help elucidate the broader evolutionary impact of TEs and their role in shaping fruit diversity.

## METHODS

### Transposable element identification and classification

The genome sequence of *Fragaria vesca* and annotation files (v4.0.a2) were downloaded from the Phytozome web portal at https://phytozome-next.jgi.doe.gov/ [116]. TEs were detected and preliminarily classified using the EDTA v2.1.0 [48], executed as follows: `EDTA.pl --genome $genome --exclude $annotation_bed --anno 1`. where `$genome` is the *F. vesca* reference genome, and `$annotation_bed` is the assembly annotation converted to BED format from the original GFF3 file using the *gff2bed* tool from BEDOPS v2.4.41 [117]. EDTA produces a filtered non-redundant TE library for annotation of structurally intact and fragmented elements. TEs are classified into several non-overlapping categories, including LTR (referring to LTR retrotransposons), non-LTR (including SINEs and LINEs), TIR (referring to DNA transposons with TIRs, including MITEs), Helitron, and non-TE repeat sequences. TEs identified by EDTA were reclassified with DeepTE [49] that uses convolutional neural networks and detects domains inside TEs to correct false classification. Within Class II, DeepTE allows for the subclassification of MITEs and non-MITEs (DNA TEs). DeepTE was run with the following command: `DeepTE.py -d /path/intermed_files -o/path/results -i $TE_lib -sp P -m_dir /path/Plants_model`. Where `$TE_lib` is the FASTA file with the TE sequences detected by EDTA, the parameter `-sp P` specifies a plant organism, and the `-m_dir` flag indicates the path to the training model. A custom bash script was then used to merge the GFF3 EDTA output files with the DeepTE classification (TE order and superfamily). For TEs categorized as “unknown” by DeepTE, the original EDTA classification was preserved, translating it into the classification used in this work (Figure 1B). The consensus TE library was then curated using TEtrimmer [118], a tool designed to automate the manual curation of TEs, which performs automated consensus sequence curation, clusters related elements, and removes duplicates. This step markedly improves TE library quality and facilitates more accurate repeat masking and classification. TEtrimmer was run with default parameters and the `--classify_all` option. Then, the consensus library file after de-duplication step was used as input for RepeatMasker to finally annotate the curated TE library.

Two GFF3 files were generated with the genomic coordinates and assigned TE class/superfamily. One includes all TEs (including fragments), while the other contains only structurally intact TEs, that is, those containing all domains corresponding to their superfamily. TE-derived IRs were classified by grouping sequences based on their sequence identity, using the CD-HIT-EST module of CD-HIT v4.8.1 [119], as previously reported for *Arabidopsis thaliana* [63], running the following command: `cd-hit-est -i $fa -o $output -c 0.8 -A 0.8 -T 2 -n 5`. A minimum sequence identity threshold of 80% (`-c`), a minimum alignment coverage of 80% (`-A`), and a word size of 5 (`-n`) were set. Next, DeepTE was run as previously described for TE annotation, using the representative sequences from each cluster to classify the IRs. When IRs overlapped with annotated TEs but were not initially classified as Class II, they were reclassified based on the overlapping element’s category using a custom bash script. The presence of transposase-associated domains in Class II elements was evaluated using the DeepTE_domains.py module, which detects conserved protein domains in TE sequences, applied to both Class II consensus sequences and genome-wide annotated copies.

### IR and MITE analysis

Inverted repeats were identified in the *F. vesca* v4 genome using *einverted* from the EMBOSS program suite v6.6.0.0 [68]. The following parameters were used: maximum repeat of 700 bp, a gap penalty of 8, a minimum score threshold of 150, a match score of 3, and a mismatch score of −4. A custom Perl script was used to parse the output and generate a GFF3 file with the IR annotation. To classify IRs as MITE-independent elements, we used bedtools intersect v2.26.0 [59], discarding elements that exhibited more than 20% overlap between MITEs and IRs.

To assess the conservation of the MITE Mutator, the Tripal MegaSearch tool from the Genome Database for Rosaceae (GDR) website (https://www.rosaceae.org/) was first used to identify orthologous genes in the strawberry species *Fragaria chiloensis* (KIB CAS genome assembly) and *Fragaria × ananassa* (Camarosa genome assembly). A 3,000 bp sequence upstream of each gene was retrieved from JBrowse (GDR) for the respective genomes and locally aligned with the *F. vesca* MITE Mutator sequence using EMBOSS Water (https://www.ebi.ac.uk/jdispatcher/psa/emboss_water), which employs the Smith-Waterman algorithm for pairwise alignment.

The predicted secondary structures with minimum free energy for each MITE were generated using the RNAfold web server (http://rna.tbi.univie.ac.at/cgi-bin/RNAWebSuite/RNAfold.cgi) with default parameters.

### Plant material and growth conditions

Plants from the *F. vesca* accession ‘Reine des Vallées’ used in this study were grown and maintained in a greenhouse under natural sunlight conditions (IHSM “La Mayora”, Málaga, Spain), using a mixture of universal substrate and river sand (3:1 ratio by volume). Fruits at the immature (white) and ripe (red) stages were collected for small RNA-seq (sRNA-seq) and RNA-seq transcriptome analyses. Strawberry fruits infiltrations were performed as previously described [120]. MITE-Mutator siRNA sponge was created by fussing half of its sequence (only one of the TIR to prevent dsRNA hairpin formation) with the GFP cloning sequence.

### sRNA-seq

RNA was extracted as previously described [121] from fruit receptacles at two developmental stages, immature (white) and ripe (red). In detail, for each stage, two biological replicates were prepared; each biological replicate was composed of 30 receptacles after the removal of the achenes. RNA quality of the samples was measured on a Bioanalyzer 2100 (Agilent Technologies, Santa Clara, CA). RNA purity was checked using the NanoPhotometer® spectrophotometer (IMPLEN, CA, USA). RNA integrity was analyzed by lab-on-a-chip electrophoresis using the RNA Nano 6000 Assay Kit with the Agilent Bioanalyzer 2100 system (Agilent Technologies, CA, USA). Quantification of samples was performed with Qubit® RNA Assay Kit in the Qubit® 3.0 Flurometer (Life Technologies, CA, USA). Subsequently, 1 μg of total RNA per sample was used to generate small RNA libraries using the Truseq® Small RNA Library Prep Kit according to manufacturer’s instructions (Illumina Inc., San Diego, CA, USA). Library pool was sequenced on the Illumina Nextseq550 instrument at the University of Málaga (Spain), with 1 × 75 cycles according to the manufacturer’s protocol. Publicly available sRNA-Seq reads from *F. vesca* leaves were retrieved from GEO under accession number GSM1513956 and analyzed together with immature and ripe reads.

The small RNA reads were first cut to remove 3’ adapters using cutadapt v4.9 and their quality checked using FastQC v0.12.1 (https://www.bioinformatics.babraham.ac.uk/projects/fastqc/) and MultiQC v1.25.1 [122]. They were then mapped with ShortStack v4.1.1 [123] to *F. vesca* v4 genome using the following parameters: `--locifile` (with MITE or IR annotation files), `--nohp` (prevents search for microRNAs) and `--make_bigwigs`. The DESeq2 Bioconductor package ([124], R Core Team 2022) was used to normalize the total raw read counts from ShortStack for differential expression analysis. MITEs/IRs with 10 or more 24-nucleotide reads across their entire region in all samples were considered to have potential siRNA production. After count normalization, the expression of IR/MITE-derived siRNA was compared between leaf tissue vs. immature stage, ripe vs. immature stages, and ripe stage vs. leaf tissue, using the negative binomial test. Differentially expressed MITEs/IRs were determined by adjusting p-values with the Benjamini–Hochberg false discovery rate, and those with an absolute log2 fold change ≥ 1 and an adjusted p-value ≤ 0.05 were defined as differentially expressed elements.

### RNA-seq

Total RNA was isolated as described previously [120] using the same protocol applied to the fruit receptacles used for sRNA-seq analysis. For the RNA-seq experiment, however, three independent biological replicates were prepared per developmental stage. After validating the quality and quantity of the total RNA using Bioanalyzer 2100 (Agilent Technologies Santa Clara, CA, USA) and Qubit 2.0, libraries were prepared using Illumina’s TruSeq RNA Sample Prep Kit v2. Pools of libraries were sequenced by paired-end sequencing (100×2) in an Illumina HiSeq 2500 sequencer by Sistemas Genómicos (Valencia, Spain). Also, transcriptome data for leaf tissue (GEO: GSM3243286 and GSM3243287), fruits treated with 5-AZA and its control (GEO: GSE151534) produced by previous studies were used [82, 125]. The analysis started by quality trimming and filtering the raw reads with Trimmomatic v0.39 [126], the reads were then aligned to the *F. vesca* v4 reference genome using STAR v2.7.11b [127] which was guided by the gene and exon annotation from *F. vesca* v4.0.a2. *Sort* and *bamCoverage* tools from Samtools v1.21 [128] were then used to sort BAM files by genomic coordinates and generate normalized coverage tracks for visualization and further analysis, respectively. Read quality before and after trimming was analyzed with FastQC v0.12.1 (https://www.bioinformatics.babraham.ac.uk/projects/fastqc/) and, together with mapping efficiency, they were summarized with MultiQC v1.25.1 [122]. Read counts on each gene were then calculated with featureCounts v2.0.8 [129]. This pipeline was run with the aid of the Snakemake workflow engine [130]. Differential expression analysis was performed by the DESeq2 package [124], filtering out genes with counts below 10 in all samples. Genes were then annotated using the information (BLAST, InterPro, GO term, GO accession) provided by the GDR database (https://www.rosaceae.org/).

### BS-seq

Publicly available Bisulfite-Seq data [125] from *F. vesca* leaf tissue, immature receptacle and ripe receptacle were retrieved from GEO under accession numbers GSM3243379, GSM3243380 and GSM3243381, respectively. Although it is not specified whether tissues were collected from the same individual plants, the observed methylation trends, including global hypomethylation and consistent DMRs across replicates, support their reliability for integrative methylome analysis. The EpiDiverse WGBS v1.0 pipeline for bisulfite read trimming, mapping and methylation calling was implemented (https://github.com/EpiDiverse/wgbs), which was specifically designed for non-model plant species [131]. For each sample, bedGraph files corresponding to each sequence context (CpG, CHG, CHH) were obtained. Then *bedGraphToBigWig* v.2.10 from ENCONDE (https://www.encodeproject.org/software/bedgraphtobigwig/) was used to generate coverage tracks for visualization and plotting.

### 3C assay and RT–qPCR

The 3C assay was conducted following a published procedure [66]. Approximately 2 g of tissue was collected per sample and crosslinked with 1% formaldehyde. Crosslinking was stopped by adding glycine to a final concentration of 0.125 M, after which nuclei were isolated. To detect FvAAT1 loops DNA was digested with XhoI (Thermo Fisher Scientific). DNA fragments were ligated with 25 U of high concentrated T4 DNA ligase (Thermo Fisher Scientific) at 22 °C for 5 h in a 2-ml volume. Subsequent steps included reverse crosslinking, proteinase K treatment (Qiagen) and DNA purification with the phenol–chloroform. To measure interaction frequency, qPCR was performed, and results were analyzed with the 2−ΔΔCt formula using ACT2 as a control. The final qPCR products were subjected to agarose gel electrophoresis (2%) and bands corresponding to the expected size were purified and sequenced using the Sanger method to confirm ligation products in the case of 3C experiments. For RT–qPCR, 500 ng of total RNA underwent RT using the RevertAid RT kit (Thermo Fisher Scientific) after treatment with DNase (Thermo Fisher Scientific). Subsequently, qPCR was conducted using SYBR green (Thermo Fisher Scientific Maxima SYBR Green qPCR Master Mix (2×)). Relative expression levels were determined using the 2−ΔΔCt method with *FvCHP1* as the housekeeping gene control.

### Identification of TE Polymorphisms

A set of 217 resequenced woodland strawberry genomes from a European germplasm collection (PRJNA1018297), representing the species’ continental diversity, was used for the polymorphism study [44]. Detailed information on each accession’s geographic origin, read depth, and chromosomal coverage can be found in Table S6. Since the study of TE polymorphisms requires a coverage over 15X and no less than 10X [90], resequencing data was analyzed using SAMtools *coverage* v1.17 [132]. The results were processed using custom bash and R scripts. Next, the accessions were filtered based on a mean per-chromosome coverage exceeding 10X. Consequently, seven of the 217 available accessions (Alta, FIN55, NOR2, NOR-P1-10, RUS3, UK3, and UK9) were excluded from subsequent analyses, resulting in a final set of 210 accessions.

TEPID v0.10 [90] was employed for the detection of polymorphisms, using the curated non-redundant TE library, and the IR annotation. The three stages of the TEPID algorithm were executed using the commands listed below:

#### Initial Mapping Stage

`tepid-map -x $bowtie_index -y $yaha_index -p $threads -s $insert_size-n $reseqid -1 $r1 -2 $r2 -z`

Where `$bowtie_index` and `$yaha_index` indicate the paths to the reference indices generated by each program; `$insert_size` refers to the average insert size of the sequencing library, set to 250 bp based on data obtained using the *CollectInsertSizeMetrics* module from Picard v3.1.1 (https://broadinstitute.github.io/picard/); `$reseqid` represents the genome under analysis; `$r1` and `$r2` denote the forward and reverse reads, respectively; and the -z flag is used to handle compressed FASTQ files.

#### Discovery Stage

`tepid-discover -p $threads --name $reseqid --conc $conc_BAM --split $split_BAM --te $TE_bed`

Where `$reseqid` is the genome identifier; `$conc_BAM` and `$split_BAM` are the Bowtie2 [133] and YAHA [134] alignment files resulting from the mapping stage, respectively; and `$TE_bed` is the BED file with TE annotations in the format specified by TEPID.

After the discovery stage, an intermediate step involved merging the TE insertions and deletions detected in the population into a single BED file using the TEPID scripts `merge_insertions.py` and `merge_deletions.py`. Briefly, insertions of the same TE were merged if their coordinates were within ±100 bp, generating a list of accessions containing that insertion. Similarly, TE deletions were merged to create a file representing all deletions detected in the population, along with the list of accessions containing each deletion.

#### Refinement Stage

`tepid-refine -p $threads --insertions $insertions --deletions $deletions --name $reseqid --conc $conc_BAM --split $split_BAM -a $reseq_list --te $TE_bed`

Where `$insertions` and `$deletions` contain the identified insertion and deletion polymorphisms for all samples in the population; `$reseqid` is the genome identifier to run the *refine* command on; `$conc_BAM` and `$split_BAM` are the Bowtie2 and YAHA alignment files, respectively; `$reseq_list` is the list with all the genome identifiers in the population; and `$TE_bed` is the TE annotation BED file.

This step, designed to reduce false-negative calls, re-evaluates genomic regions with less stringent settings where a TE variant was identified in at least one sample of the population but not in the focal sample. It produces two types of BED files for each accession: (i) second_pass, referring to polymorphisms not detected in the discovery step but rescued after re-evaluating the region, and (ii) ambiguous, referring to TE presence/absence calls that could not be reliably determined during the refinement step. For each accession, second_pass insertions and deletions were combined with the variants discovered in step 2. The resulting final insertion and deletion files for each accession were then merged using the scripts `merge_insertions.py` and `merge_deletions.py`.

The resulting BED files were processed using custom R scripts.

To determine the genomic distribution of TE polymorphisms (Figure 6C and Figure S10) we developed a custom script using R packages GenomicRanges and GenomicFeatures [135]. Insertions and deletions were mapped against the *Fragaria vesca* v4.0.a2 reference annotation. Genomic features were defined hierarchically to resolve overlapping annotations: regions 1 kb upstream of the transcription start site (TSS) were classified as promoters (“Upstream”), followed by 1 kb downstream of the transcription termination site (TTS). Within gene bodies, determining priority was given to Exons, followed by UTRs (5’ and 3’), and finally Introns. Polymorphisms falling outside these regions were classified as distal intergenic (>1 kb or <-1 kb). To assess whether the observed distribution deviated from random expectation, we calculated the “Genomic fraction” occupied by each feature type. This was calculated by collapsing the reference genome into non-overlapping unique base pairs (unstranded) to represent the physical target size available for mutation. Enrichment or depletion was quantified using the *Log_2_* ratio of the Observed frequency to the Expected frequency (percentage of the genome occupied by a specific feature).

### Genome-Wide Association Study

GWAS was conducted on the 4395 and 4340 polymorphisms present in 116 and 154 accessions for harvests H16 and H17 [91], respectively, with a MAF ≥ 5%. The association analyses were performed by GAPIT (version 3.5) [136] using a Multi-Locus Mixed Model (MLMM) [137]. The model included the three first principal components derived from all the polymorphisms used for each group of accessions (H16 and H17) (Figure S11) as covariates to reduce false positives due to population stratification. Additionally, a SNP-based kinship matrix capturing the relationship between accessions was included as random effects. Biallelic variants with a minor allele frequency greater than 0.01 were retained for kinship estimation using the VanRaden method [138]. The Bonferroni correction was used to establish a threshold of significance for all the associations.

### Data processing and visualization

To calculate the number of genes near each type of TE and differentially expressed MITEs/IRs, the tool *slop* and *intersect* from bedtools v2.26.0 [59] were used for distances of 500, 1000, and 3000 bp both upstream and downstream of each element. Additionally, *intersect* was used to count elements located within coding regions, utilizing different annotation files for each feature.

To evaluate whether MITEs or IRs are preferentially associated with gene expression changes relative to other genomic contexts, genes were classified into mutually exclusive categories based on their proximity (≤ 3 kb) to: (i) MITEs or IRs, (ii) other TE classes, or (iii) no annotated TE (TE-free regions). Each gene was assigned to a single category based on genomic overlap identified using bedtools intersect. For each comparison, genes were further classified as DE or non-DE using the list of DE genes identified from the RNA-seq analysis comparing mature versus immature fruit stages. Contingency tables were constructed, and statistical significance was assessed using Fisher’s exact test. Odds ratios (OR) and p-values were used to estimate the relative likelihood of differential expression across genomic contexts.

In order to evaluate the global methylation and siRNA level, metagene analysis was performed using deepTools v3.5.5 [125]. First, *computeMatrix* was executed to generate read abundance from all samples over gene, TE, MITE and IR regions. This matrix was then used to create, using *plotProfile*, a metagene profile from 2 kb upstream of the Transcription Start Site (TSS) to 2 kb downstream of the Transcription End Site (TES).

Moreover, circos ideograms were created using Circos software vv0.69-6 [126], displaying only the assembled chromosomes due to their higher density of gene and transposon annotations. Also, JBrowse2 desktop v2.18.0 [127] was used to visualize gene and MITE/IR annotations, along with RNA-seq and BS-seq data, at specific *loci* in the *F. vesca* v4 genome. Principal Component Analysis (PCA) was performed using the R package prcomp [139]. Data obtained from the different analyses of TE annotation and from the sequencing experiments were further processed, statistically analyzed, and plotted in R using a diversity of packages, including dplyr (https://github.com/tidyverse/dplyr), tidyr (https://github.com/tidyverse/tidyr), scales (https://github.com/r-lib/scales), forcats (https://github.com/tidyverse/forcats), ggplot2, RColorBrewer (https://CRAN.R-project.org/package=RColorBrewer), MetBrewer (https://github.com/BlakeRMills/MetBrewer), rtracklayer [140], and ggman (https://github.com/drveera/ggman).

## Ethics approval and consent to participate

Not applicable.

## Consent for publication

Not applicable.

## Availability of data and materials

The datasets generated during the current study are available from European Nucleotide Archive (ENA) under the accession number PRJEB86847. The curated TE, MITE, and IR libraries, as well as the code for TE polymorphism and GWAS analyses, are available at: https://github.com/spriego/PriegoCubero_Tolley_et_al_Fvesca

## Authors’ contributions

**R.T.** carried out TE identification and classification. **I.T.** conducted MITEs and IRs bioinformatic analysis. **S.P.C.** performed TE polymorphism identification and GWAS analysis. **J.L.G**. and **C.Z** conducted all the validations of the MITE-triggered effects over *FvAAT1*. **C.M.P.** collected the samples and performed sRNA-sequencing and RNA-seq experiments. **T.H.**, and **T.T.** provided genomic data of *F. vesca* accessions (bam files). **I.T.**, **I.T., C.M.P.**, and **P.A.M.** conceived the study. **P.A.M.** secured project funding. **I.T.** and **P.A.M.** supervised the work and wrote the manuscript with the help of all authors.

## Acknowledgements

We would like to acknowledge the Sequencing Unit from the University of Malaga for the support they have provided for the small RNA-seq analysis. We thankfully acknowledge the computer resources (Picasso Supercomputer), technical expertise and assistance provided by the SCBI (Supercomputing and Bioinformatics) center of the University of Malaga. This work was supported by grants from the Spanish Ministry of Science and Innovation (PID2022-137037NB-I00 and CNS2023-145312) to P.A.M.

## SUPPLEMENTARY MATERIAL

**Table S1.** Size distribution of Transposable Elements by Class annotated in *Fragaria vesca*, including both fragmented and intact TEs.

**Table S2.** ShortStack analysis of sRNA-seq data from *Fragaria vesca* leaves, immature (white stage), and ripe (red stage) fruits using annotated loci files for MITEs and IRs.

**Table S3.** Differential expression analysis of 24-nt siRNA-producing MITEs and adjacent genes (< 3 kb) in *Fragaria vesca*. Genomic coordinates, functional annotation and expression changes between leaf tissue, and immature and ripe stages are shown. Gene IDs associated with key roles in fruit development and metabolism are indicated in bold.

**Table S4.** Differential expression analysis of 24-nt siRNA-producing IRs and adjacent genes (< 3 kb) in *Fragaria vesca*. Genomic coordinates, functional annotation and expression changes between leaf tissue, and immature and ripe stages are shown. Gene IDs associated with key roles in fruit development and metabolism are indicated in bold.

**Table S5.** Differential gene expression analysis of all *F. vesca* genes across conditions: leaf tissue vs. immature stage, ripe stage vs. leaf tissue, ripe vs. immature stages, and AZA-treated vs. control strawberries.

**Table S6.** Resequenced genomes of *Fragaria vesca* analyzed in the present study, including accession identifiers, country and origin coordinates, read counts, and average chromosome coverage (excluding unassembled contigs). Accessions in italics were excluded from analysis due to low coverage; accessions in bold were included in the GWAS study.

**Table S7**. Summary of significant genome-wide associations between Transposable Element Polymorphisms (TEPs) and agronomic and volatilome traits across two harvest seasons (H16 and H17) in *Fragaria vesca*. Associations described in the Results section are highlighted in bold.

**Figure S1. Annotation and classification of transposable elements (TEs) in the *Fragaria vesca* v4.0.a2 reference genome. (A)** Representation of the pipeline used in this work for transposon annotation. Software employed are highlighted in blue boxes, while input and output files are shown in pink boxes, with their formats in parentheses. **(B)** Distribution of TE superfamilies as annotated by EDTA (left), by DeepTE (center), and by TEtrimmer (right) for the *F. vesca* v4.0 genome.

**Figure S2. The average length of intact TEs and all annotated TEs in *F. vesca* genome (including fragmented TEs).**

**Figure S3. Distribution of intact TE superfamilies in the *Fragaria vesca* v4.0.a2 annotation based on their distance from a gene.** Categories include: within a gene (‘intragenic’), within 1 kb (‘1000’), between 1 and 3 kb (‘3000’), and more than 3 kb (‘>3000’). Left: Total counts by superfamily. Right: Proportion by superfamily.

**Figure S4. (A)** Genome distribution of IRs and MITEs. **(B)** Proportion of IRs classified as ‘Other’, excluding MITE-type elements. Values are shown as percentages relative to the total number of IRs in the ‘Other’ category (16.1%).

**Figure S5. (A)** Metagene profile of 24-nt siRNAs mapping to MITEs and IRs located at different ranges of distance to the closest protein-coding gene, in leaf (green), and immature (gray) and ripe fruits (pink). Plots show MITEs/IRs scaled from the start to the end plus 2,000 bp to each side. sRNA-seq replicates are plotted together. **(B)** Metagene profile of CpG, CHG, and CHH DNA methylation at MITEs/IRs located at different ranges of distance to the closest protein-coding gene, in leaf (green), and immature (gray) and ripe fruits (pink). Plots show MITEs/IRs scaled from the start to the end plus 2,000 bp to each side. Individual BS-seq replicates are plotted.

**Figure S6.** Genomic region of the *F. vesc*a v4 assembly containing the MITE Mutator locus near FvH4_7g18550 and FvH4_7g18570 genes. IRs and MITEs identified in this region are also displayed.

**Figure S7. Conservation of the MITE Mutator from *Fragaria vesca* in the orthologous genomic regions of commercial strawberry species *Fragaria chiloensis* and *Fragaria × ananassa*. (A)** Local alignment of *F. vesca* MITE Mutator with the MITE located near the FvH4_7g18570 ortholog in *F. chiloensis* (left) and *F. × ananassa* (right). **(B)** Minimum free energy (MFE) RNA secondary structure predicted by RNAfold server of each MITE. The color scale shows base-pairing probability 0 (purple) to 1 (red).

**Figure S8.** Region of the *F. vesca* v4.0.a2 reference genome containing the FvH4_6g46420 **(A)**, FvH4_2g30230 **(B),** and FvH4_5g23180 **(C)** loci displaying the IR and MITE elements annotated in the region and expression and epigenetic profiles. (I) 24-nt siRNAs mapping to the genomic regions as determined by sRNA sequencing of leaf (green), immature (gray) and ripe (pink) tissues. Replicates are plotted together. (II) Expression of each gene in leaf (green), immature (gray) and ripe (pink) tissues as well as strawberries treated with 5-azacytidine (5-AZA) and its control, measured by RNA-seq. (III) Cytosine DNA methylation in CG, CHG, and CHH contexts in leaf (green), immature (gray) and ripe (pink) fruits. The average of individual BS-seq replicates is plotted for each condition.

**Figure S9. (A)** Representation of the pipeline used in this study. The main inputs are shown at the top, while the main outputs are placed at the bottom. File formats are indicated in parentheses. **(B)** Representation of the TEPID pipeline used for identifying transposable element insertion polymorphisms (TIPs) and absence polymorphisms (TAPs).

**Figure S10. Enrichment analysis of TE insertion and deletion polymorphisms across genomic features in *Fragaria vesca* v4.0.a2. (A)** Genome-wide enrichment of all polymorphisms, insertions, and deletions across annotated genomic features. **(B)** Enrichment profiles of TE-associated polymorphisms across major TE superfamilies. Values are shown as log2(observed/expected), where 0 indicates a random distribution relative to genomic fraction. Positive values (red) indicate enrichment, whereas negative values (blue) indicate depletion. Bubble size reflects the absolute magnitude of enrichment or depletion.

**Figure S11. Principal Component Analysis (PCA) of the subset of *Fragaria vesca* accessions used for GWAS analysis.** Each point represents a single accession, with colors indicating their respective country of origin.

**Figure S12. Genomic region of the *F. vesca* v4.0.a2 reference genome harboring selected TEP polymorphisms on chromosome Fvb3 associated with variation in: (A)** 6-methyl-5-hepten-2-one levels. (I) Coverage of mapped reads (light green) and split reads (dark green) in some accessions carrying the polimorphism (from top to bottom: ES14, ES-13, UK11, and IT15). (II) Coverage of mapped reads (light blue) and split reads (dark blue) in non-carriers accesions (from top to bottom: FIN10, FIN12, FIN6, and FIN51). **(B)** γ-decalactone levels. (I) Coverage of mapped reads (light green) and split reads (dark green) in some accessions carrying the polimorphism (from top to bottom: DK1, ES14, IT6, and IT15). (II) Coverage of mapped reads (light blue) and split reads (dark blue) in non-carriers accesions (from top to bottom: FR1, FIN25, FIN51, and NOR131-2).

**Figure S13. Addition TE polymorphisms linked to volatile compound variation in a collection of *F. vesca* accessions. (A)** Manhattan plot of TEP-GWAS for methyl 2-aminobenzoate in harvest season H16. The dotted gray line marks the significance threshold determined by the Bonferroni correction, and the significantly associated polymorphism is highlighted. **(E)** Observed and expected distribution of *p* values for TEP-GWAS for methyl 2-aminobenzoate. **(F)** Violin plot showing the relative abundance of methyl 2-aminobenzoate in accessions carrying the MITE Mutator deletion (+, coral) versus non-carriers (-, steel blue). Horizontal lines in (-) and (+) indicate the mean. **(D)** Manhattan plot of TEP-GWAS for γ-decalactone across both harvest seasons (H16 and H17, represented by circles and triangles, respectively). The dotted gray line marks the significance threshold determined by the Bonferroni correction, and the significantly associated polymorphisms are highlighted. **(E)** Observed and expected distribution of *p* values for TEP-GWAS for γ-decalactone. **(F)** Violin plot showing the relative abundance of γ-decalactone in accessions carrying the MITE deletion (+, coral) versus non-carriers (-, steel blue). Horizontal lines in (-) and (+) indicate the mean.

