## Supplemental information for "Naturally occurring variation in gene-associated transposable elements impacts gene expression and phenotypic diversity in woodland strawberry"

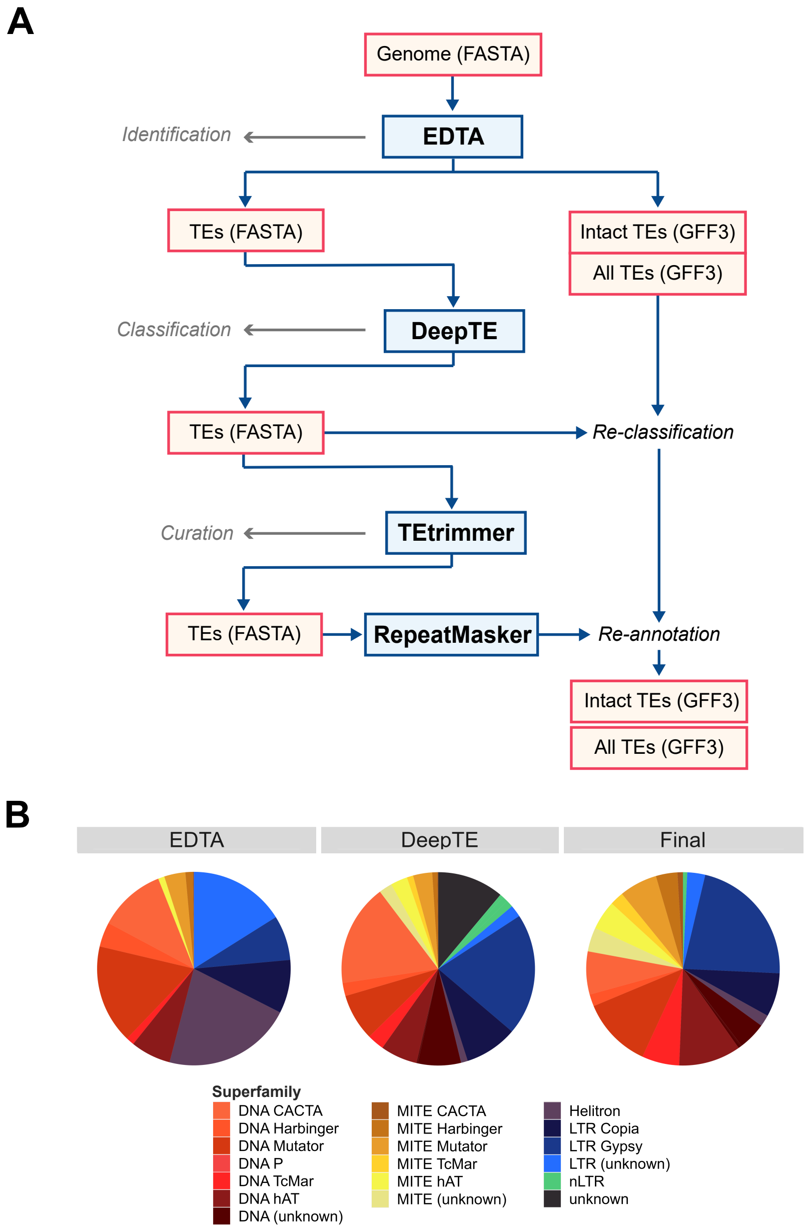


**Figure S1. Annotation and classification of transposable elements (TEs) in the *Fragaria vesca* v4.0.a2 reference genome. (A)** Representation of the pipeline used in this work for transposon annotation. Software employed are highlighted in blue boxes, while input and output files are shown in pink boxes, with their formats in parentheses. **(B)** Distribution of TE superfamilies as annotated by EDTA (left), by DeepTE (center), and by TEtrimmer (right) for the *F. vesca* v4.0 genome.

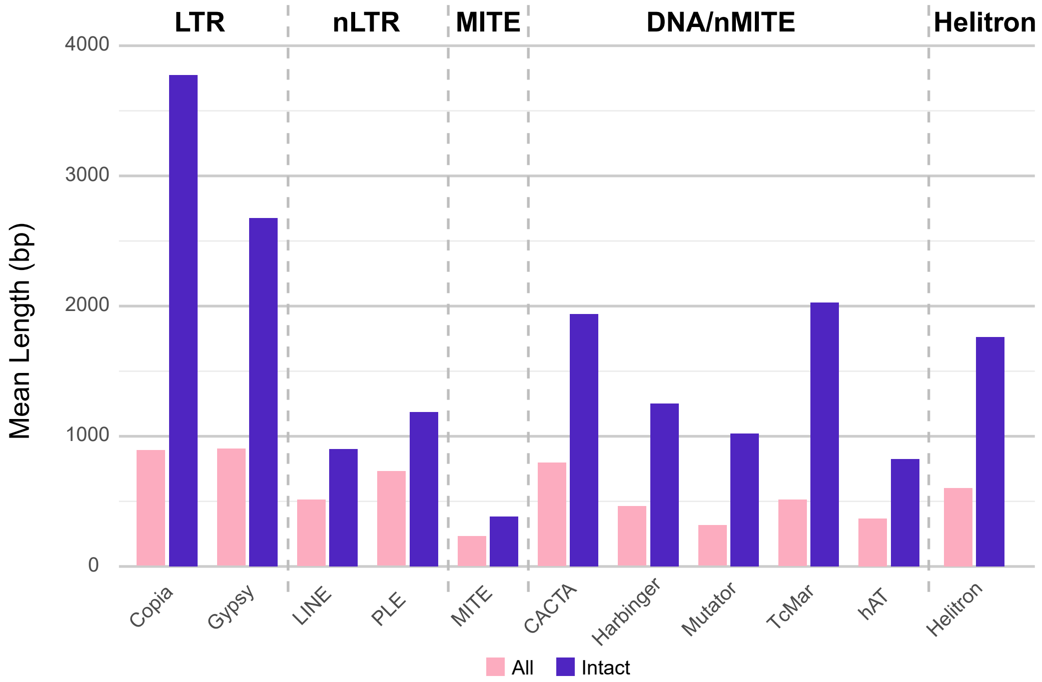


**Figure S2. The average length of intact TEs and all annotated TEs in *F. vesca* genome (including fragmented TEs).**

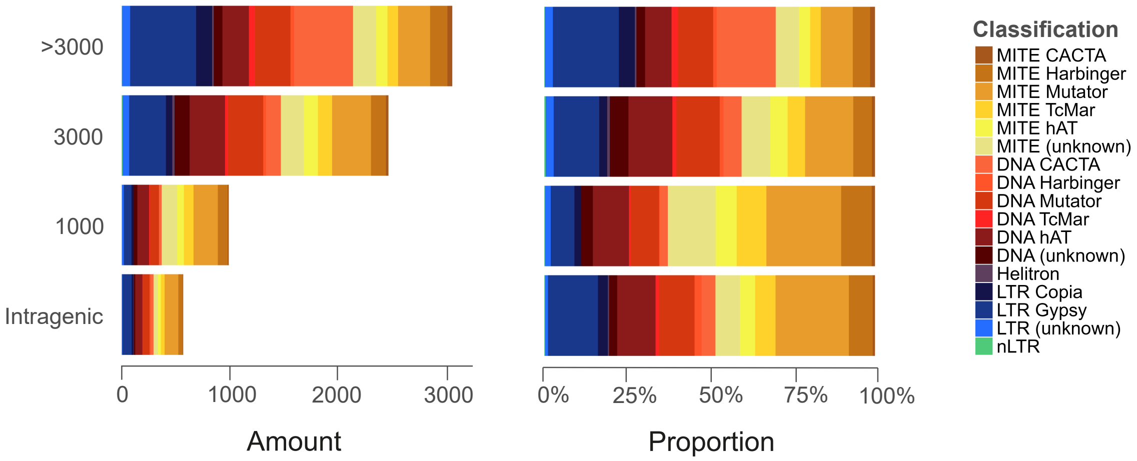


**Figure S3. Distribution of intact TE superfamilies in the *Fragaria vesca* v4.0.a2 annotation based on their distance from a gene**. Categories include: within a gene ('intragenic'), within 1 kb ('1000'), between 1 and 3 kb ('3000'), and more than 3 kb ('>3000'). Left: Total counts by superfamily. Right: Proportion by superfamily.

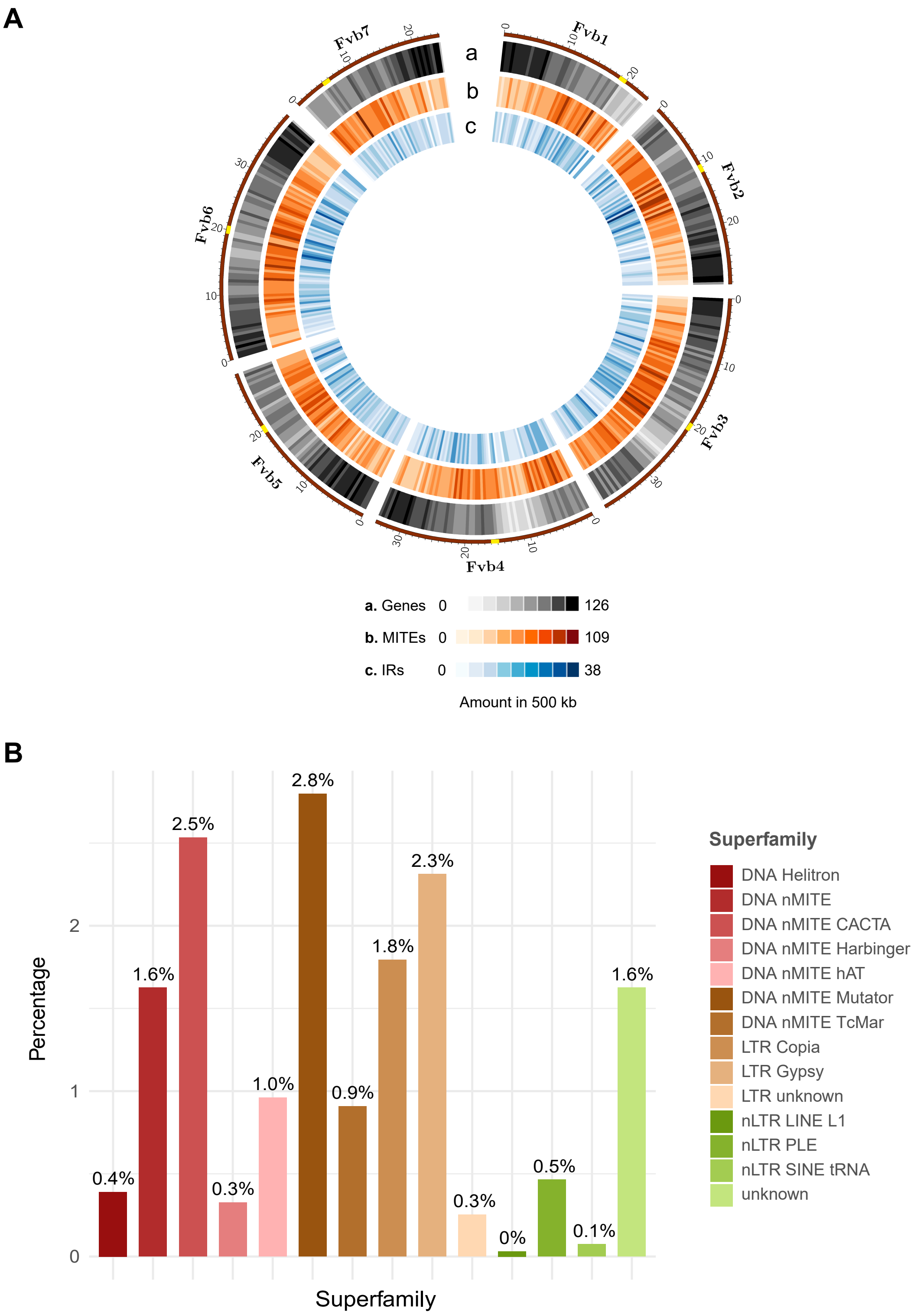


**Figure S4. (A)** Genome distribution of IRs and MITEs. **(B)** Proportion of IRs classified as ‘Other’, excluding MITE-type elements. Values are shown as percentages relative to the total number of IRs in the ‘Other’ category (16.1%).

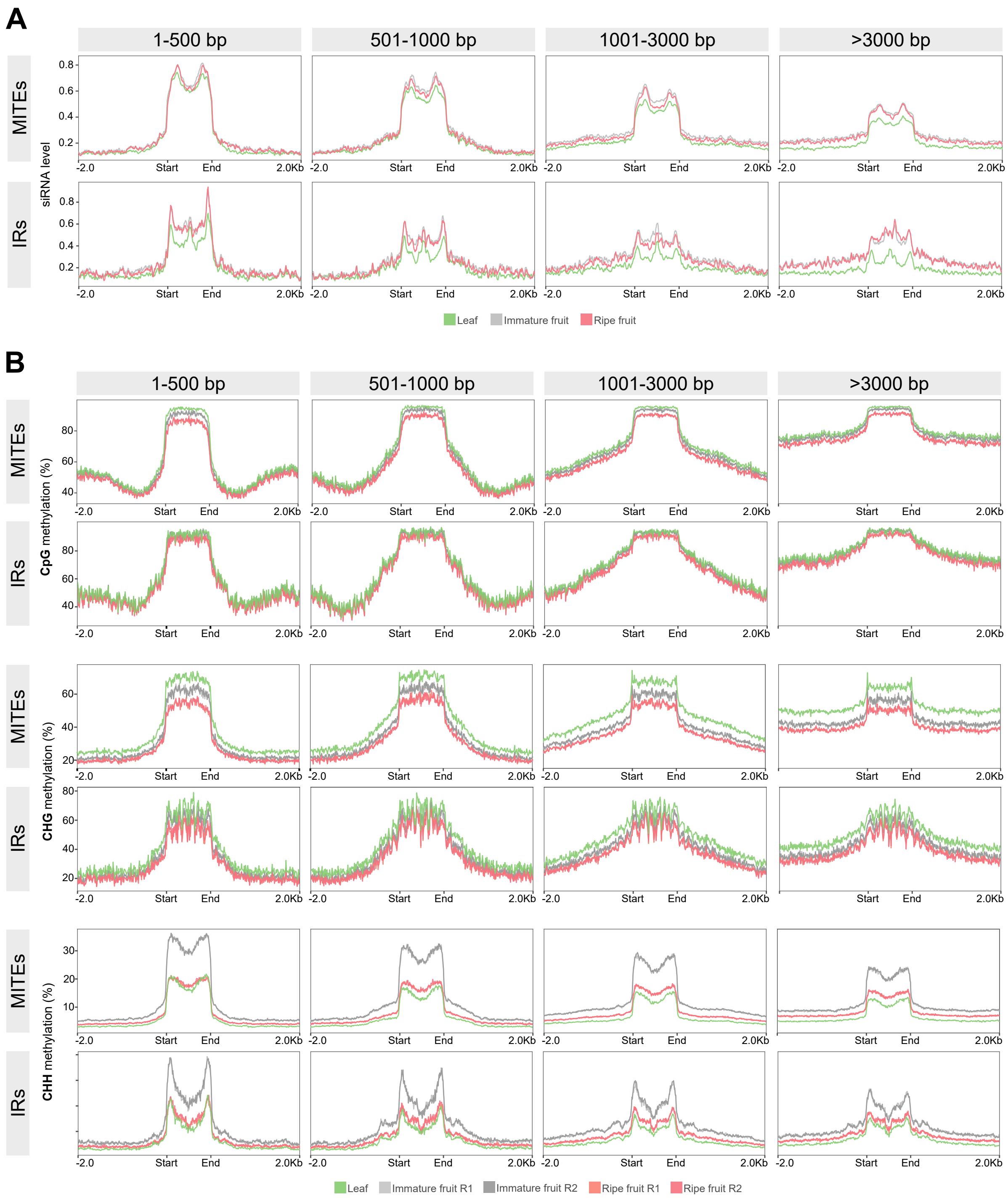


**Figure S5. (A)** Metagene profile of 24-nt siRNAs mapping to MITEs and IRs located at different ranges of distance to the closest protein-coding gene, in leaf (green), and immature (gray) and ripe fruits (pink). Plots show MITEs/IRs scaled from the start to the end plus 2,000 bp to each side. sRNA-seq replicates are plotted together. **(B)** Metagene profile of CpG, CHG, and CHH DNA methylation at MITEs/IRs located at different ranges of distance to the closest protein-coding gene, in leaf (green), and immature (gray) and ripe fruits (pink). Plots show

MITEs/IRs scaled from the start to the end plus 2,000 bp to each side. Individual BS-seq replicates are plotted.

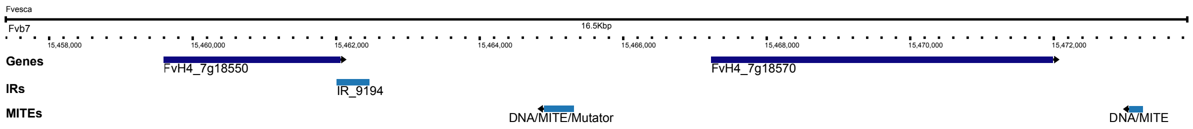


**Figure S6.** Genomic region of the *F. vesca* v4 assembly containing the MITE Mutator locus near FvH4_7g18550 and FvH4_7g18570 genes. IRs and MITEs identified in this region are also displayed.

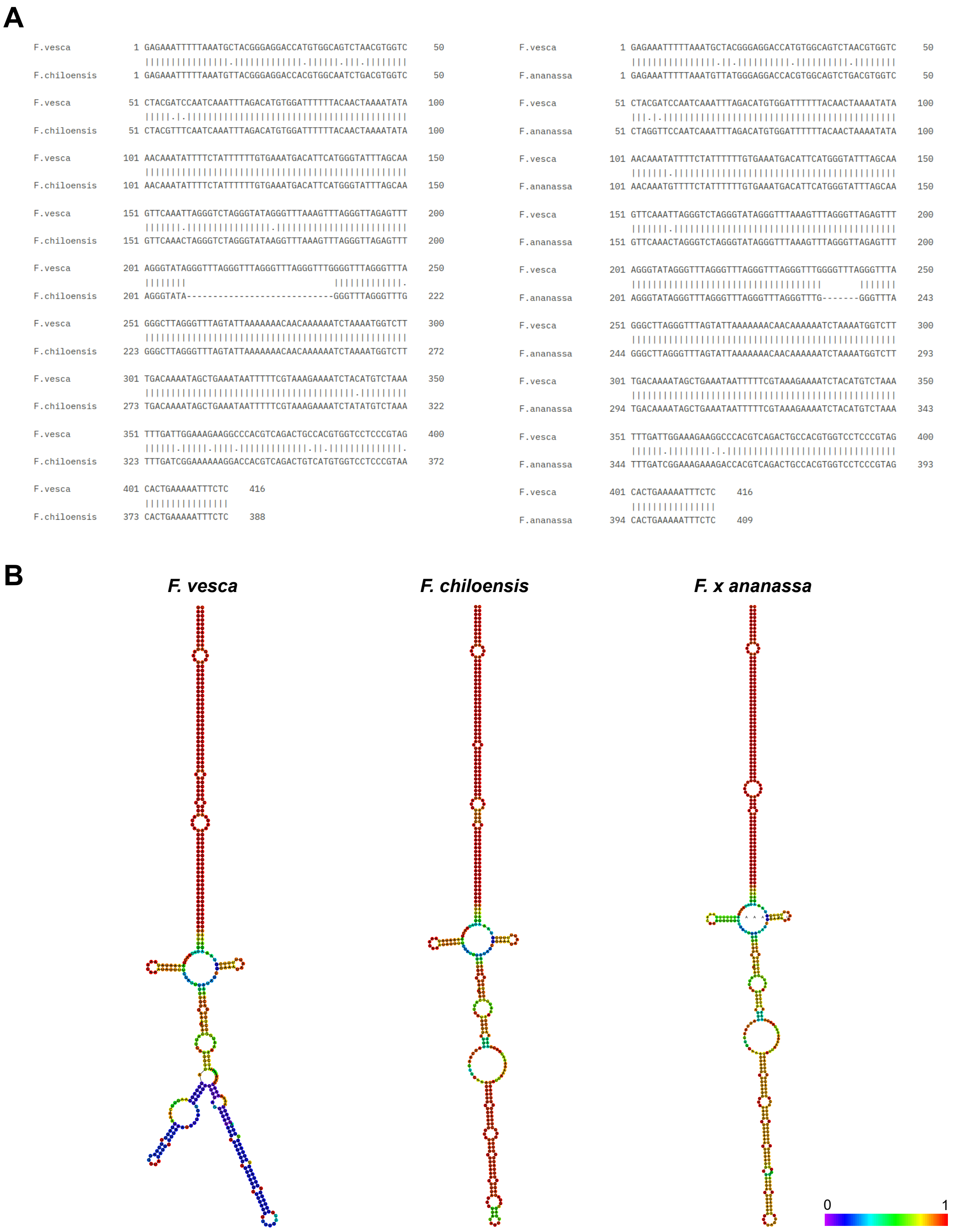


**Figure S7. Conservation of the MITE Mutator from *Fragaria vesca* in the orthologous genomic regions of commercial strawberry species *Fragaria chiloensis* and *Fragaria* × *ananassa*. (A)** Local alignment of *F. vesca* MITE Mutator with the MITE located near the FvH4_7g18570 ortholog in *F. chiloensis* (left) and *F.* × *ananassa* (right). **(B)** Minimum free energy (MFE) RNA secondary structure predicted by RNAfold server of each MITE. The color scale shows base-pairing probability 0 (purple) to 1 (red).

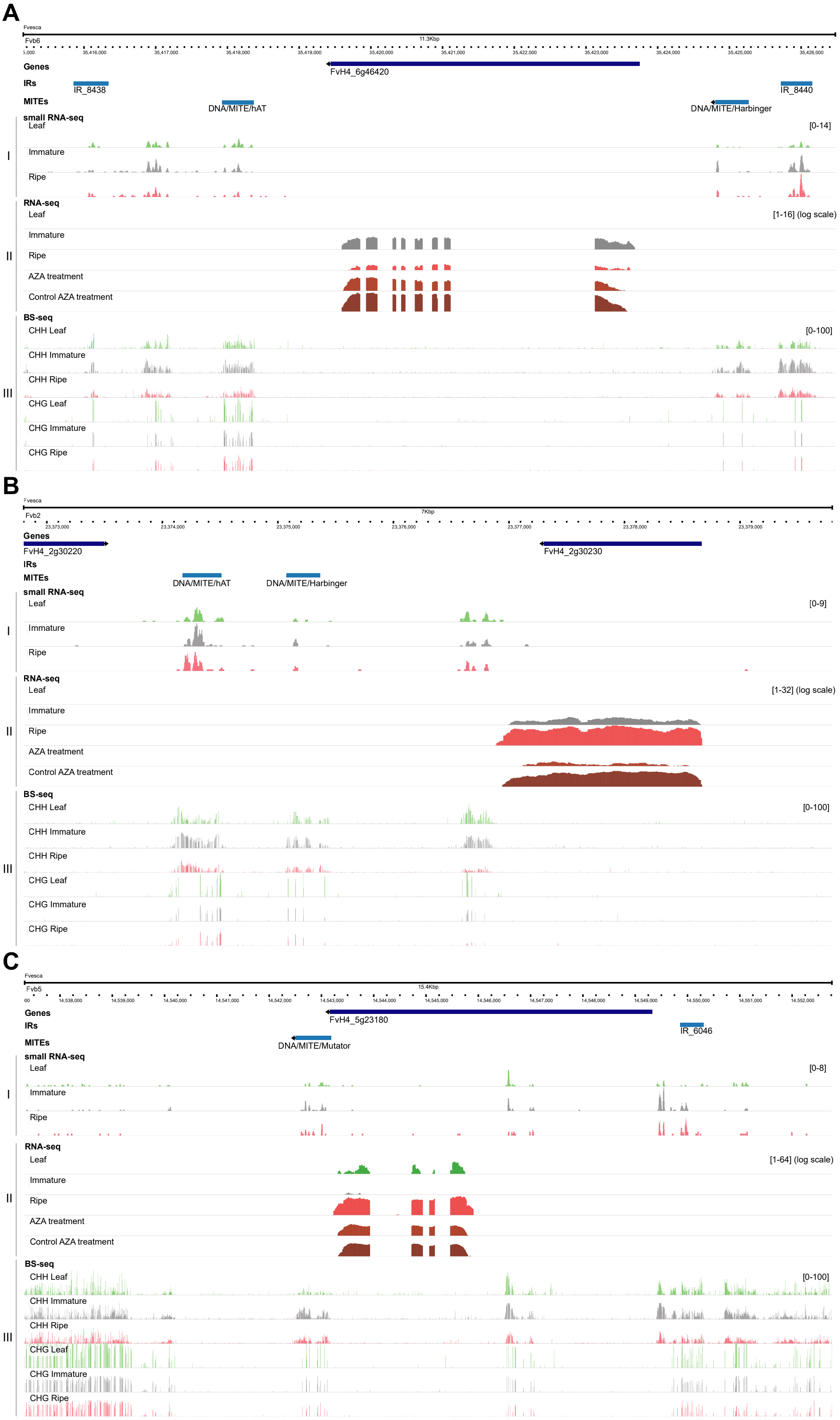
A
Fvesca
Fvb6
11.3Kbp
Genes
FvH4_6g46420
IRs
IR_8438
IR_8440
MITEs
DNA/MITE/hAT
DNA/MITE/Harbinger
small RNA-seq
Leaf
[0-14]
Immature
Ripe
I
RNA-seq
Leaf
[1-16] (log scale)
Immature
Ripe
AZA treatment
Control AZA treatment
II
BS-seq
CHH Leaf
[0-100]
CHH Immature
CHH Ripe
CHG Leaf
CHG Immature
CHG Ripe
III
B
Fvesca
Fvb2
7Kbp
Genes
FvH4_2g30220
FvH4_2g30230
IRs
MITEs
DNA/MITE/hAT
DNA/MITE/Harbinger
small RNA-seq
Leaf
[0-9]
Immature
Ripe
I
RNA-seq
Leaf
[1-32] (log scale)
Immature
Ripe
AZA treatment
Control AZA treatment
II
BS-seq
CHH Leaf
[0-100]
CHH Immature
CHH Ripe
CHG Leaf
CHG Immature
CHG Ripe
III
C
Fvesca
Fvb5
15.4Kbp
Genes
FvH4_5g23180
IRs
IR_6046
MITEs
DNA/MITE/Mutator
small RNA-seq
Leaf
[0-8]
Immature
Ripe
I
RNA-seq
Leaf
[1-64] (log scale)
Immature
Ripe
AZA treatment
Control AZA treatment
II
BS-seq
CHH Leaf
[0-100]
CHH Immature
CHH Ripe
CHG Leaf
CHG Immature
CHG Ripe
III

**Figure S8.** Region of the *F. vesca* v4.0.a2 reference genome containing the FvH4_6g46420 **(A)**, FvH4_2g30230 **(B),** and FvH4_5g23180 **(C)** loci displaying the IR and MITE elements annotated in the region and expression and epigenetic profiles. (I) 24-nt siRNAs mapping to the genomic regions as determined by sRNA sequencing of leaf (green), immature (gray) and ripe (pink) tissues. Replicates are plotted together. (II) Expression of each gene in leaf (green), immature (gray) and ripe (pink) tissues as well as strawberries treated with 5-azacytidine (5-AZA) and its control, measured by RNA-seq. (III) Cytosine DNA methylation in CG, CHG, and CHH contexts in leaf (green), immature (gray) and ripe (pink) fruits. The average of individual BS-seq replicates is plotted for each condition.

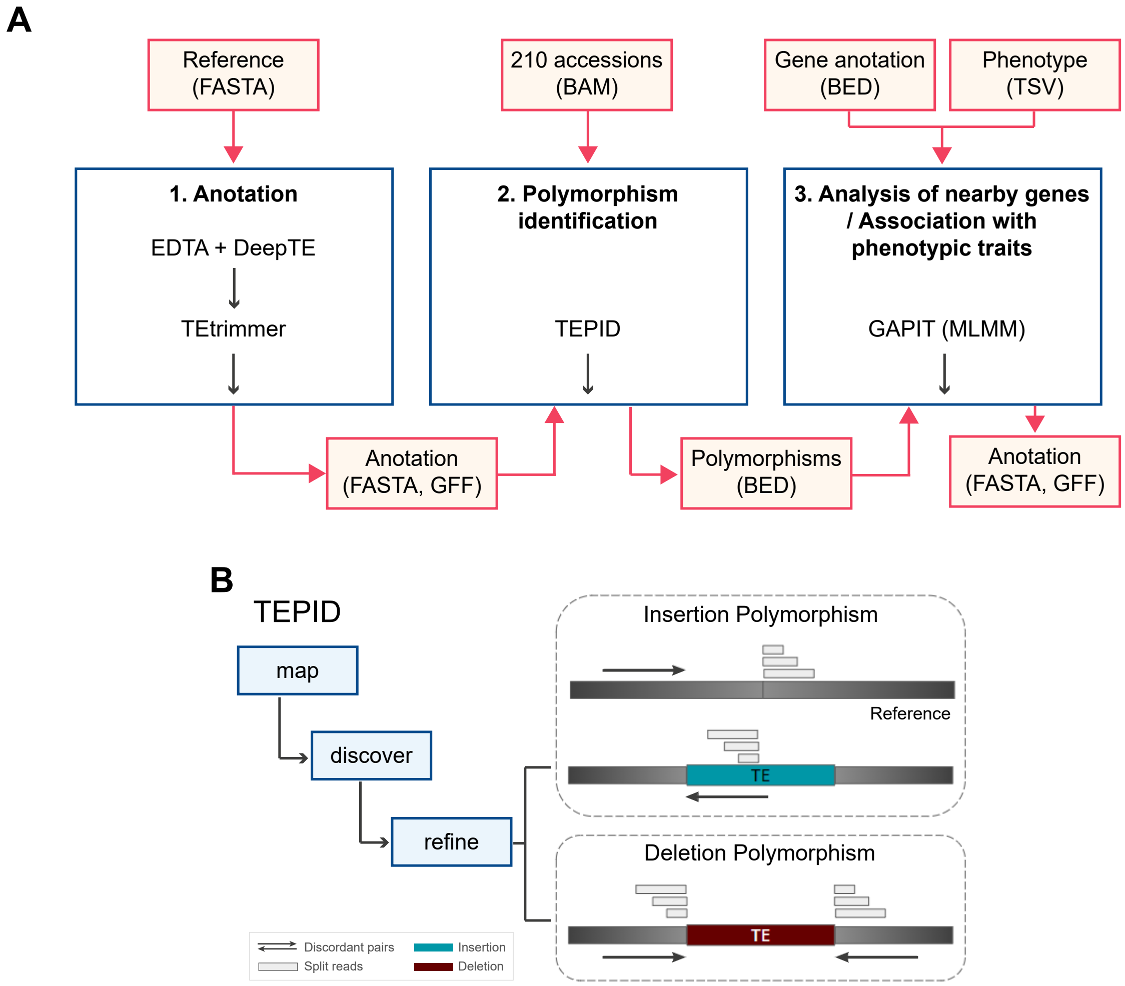


**Figure S9. (A)** Representation of the pipeline used in this study. The main inputs are shown at the top, while the main outputs are placed at the bottom. File formats are indicated in parentheses. **(B)** Representation of the TEPID pipeline used for identifying transposable element insertion polymorphisms (TIPs) and absence polymorphisms (TAPs).

A

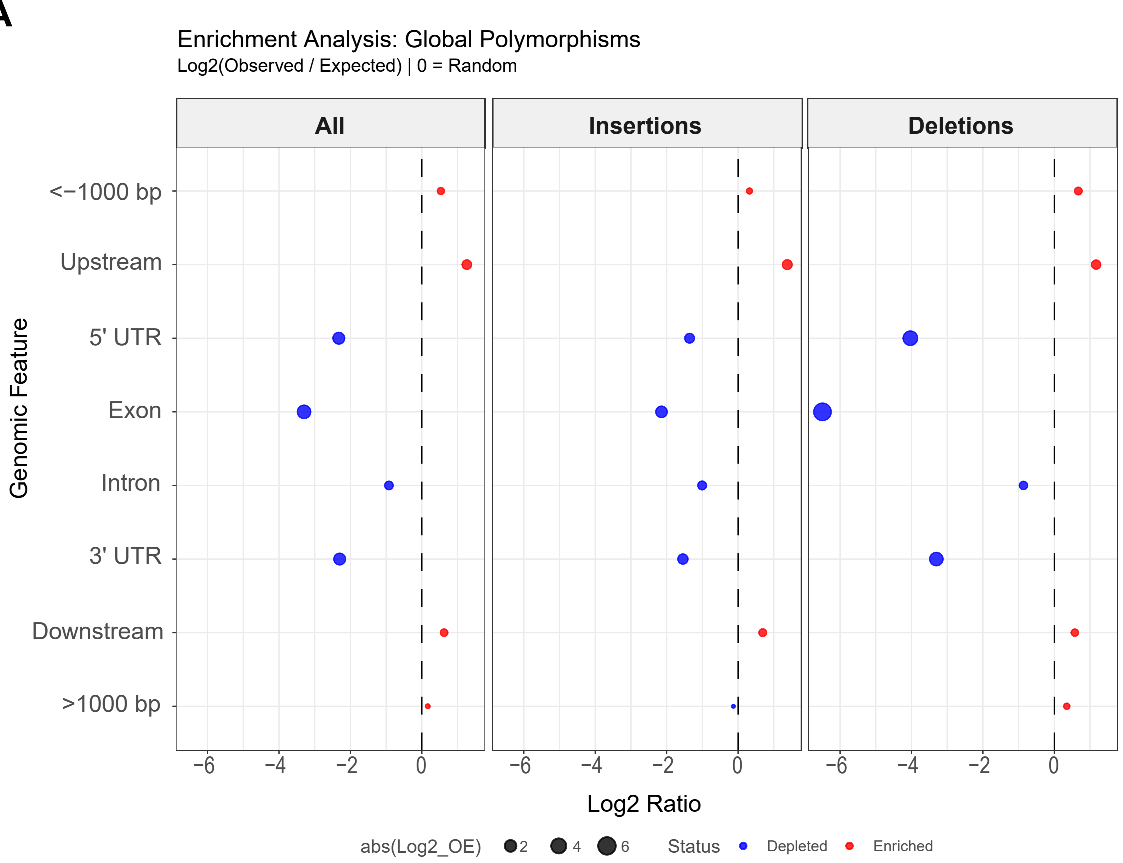
Enrichment Analysis: Global Polymorphisms
Log2(Observed / Expected) | 0 = Random
All
Insertions
Deletions
<−1000 bp
Upstream
5' UTR
Exon
Intron
3' UTR
Downstream
>1000 bp
Genomic Feature
−6
−4
−2
0
Log2 Ratio
abs(Log2_OE)
2
4
6
Status
Depleted
Enriched

B

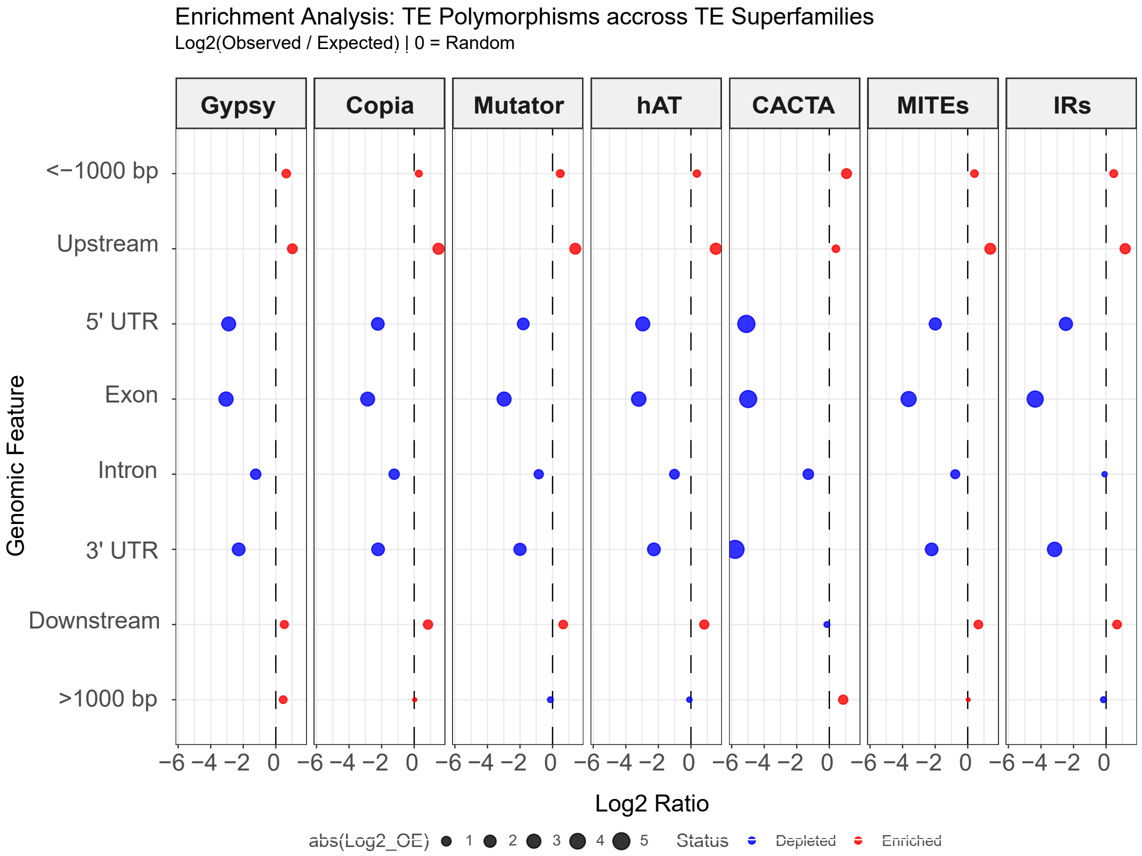
Enrichment Analysis: TE Polymorphisms accross TE Superfamilies
Log2(Observed / Expected) | 0 = Random
Gypsy
Copia
Mutator
hAT
CACTA
MITEs
IRs
<−1000 bp
Upstream
5' UTR
Exon
Intron
3' UTR
Downstream
>1000 bp
Genomic Feature
−6 −4 −2 0
Log2 Ratio
abs(Log2_OE)
1
2
3
4
5
Status
Depleted
Enriched

**Figure S10. Enrichment analysis of TE insertion and deletion polymorphisms across genomic features in *Fragaria vesca* v4.0.a2. (A)** Genome-wide enrichment of all polymorphisms, insertions, and deletions across annotated genomic features. **(B)** Enrichment profiles of TE-associated polymorphisms across major TE superfamilies. Values are shown as log2(observed/expected), where 0 indicates a random distribution relative to genomic fraction. Positive values (red) indicate enrichment, whereas negative values (blue) indicate depletion. Bubble size reflects the absolute magnitude of enrichment or depletion.

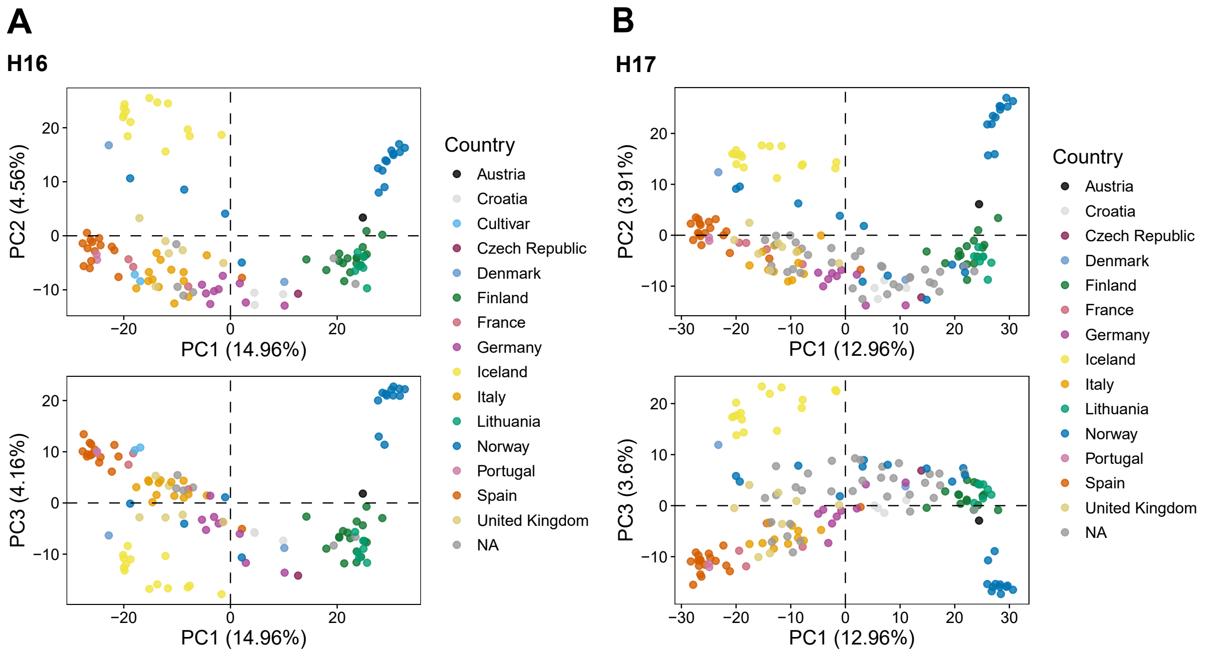


**Figure S11. Principal Component Analysis (PCA) of the subset of *Fragaria vesca* accessions used for GWAS analysis.** Each point represents a single accession, with colors indicating their respective country of origin.

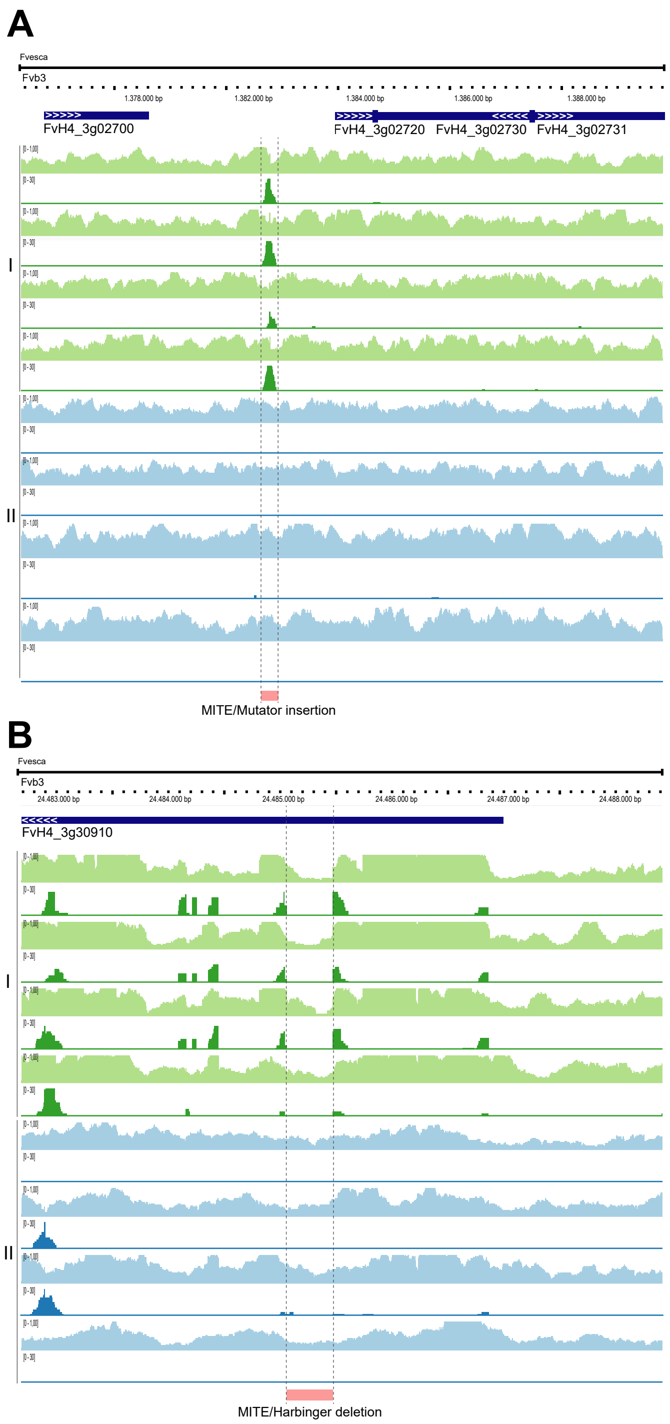


**Figure S12. Genomic region of the *F. vesca* v4.0.a2 reference genome harboring selected TEP polymorphisms on chromosome Fvb3 associated with variation in: (A)** 6-methyl-5-hepten-2-one levels. (I) Coverage of mapped reads (light green) and split reads (dark green) in some accessions carrying the polimorphism (from top to bottom: ES14, ES-13, UK11, and IT15). (II) Coverage of mapped reads (light blue) and split reads (dark blue) in non-carriers accesions (from top to bottom: FIN10, FIN12, FIN6, and FIN51). **(B)** γ-decalactone levels. (I) Coverage of mapped reads (light green) and split reads (dark green) in some accessions carrying the polimorphism (from top to bottom: DK1, ES14, IT6, and IT15). (II) Coverage of mapped reads (light blue) and split reads (dark blue) in non-carriers accesions (from top to bottom: FR1, FIN25, FIN51, and NOR131-2).

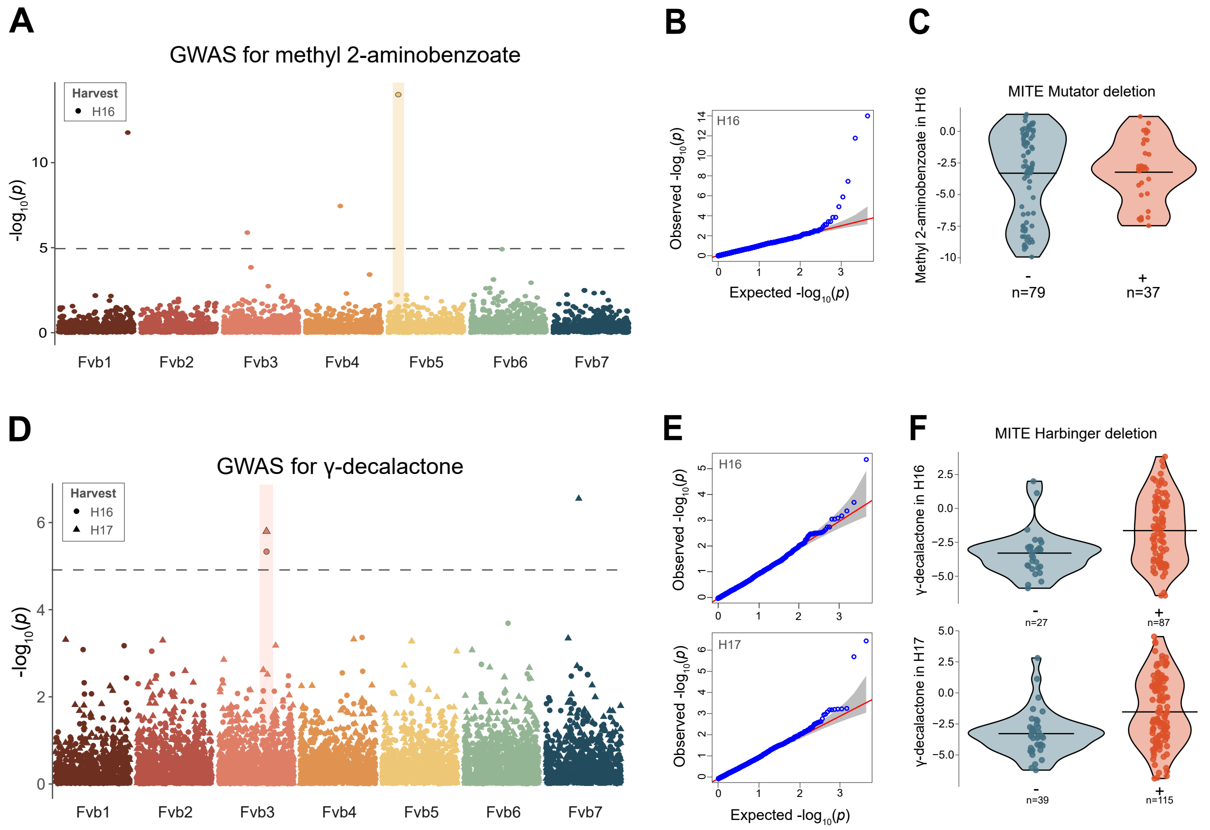


**Figure S13. Addition TE polymorphisms linked to volatile compound variation in a collection of *F. vesca* accessions. (A)** Manhattan plot of TEP-GWAS for methyl 2-aminobenzoate in harvest season H16. The dotted gray line marks the significance threshold determined by the Bonferroni correction, and the significantly associated polymorphism is highlighted. **(E)** Observed and expected distribution of *p* values for TEP-GWAS for methyl 2-aminobenzoate. **(F)** Violin plot showing the relative abundance of methyl 2-aminobenzoate in accessions carrying the MITE Mutator deletion (+, coral) versus non-carriers (-, steel blue). Horizontal lines in (-) and (+) indicate the mean. **(D)** Manhattan plot of TEP-GWAS for γ-decalactone across both harvest seasons (H16 and H17, represented by circles and triangles, respectively). The dotted gray line marks the significance threshold determined by the Bonferroni correction, and the significantly associated polymorphisms are highlighted. **(E)** Observed and expected distribution of *p* values for TEP-GWAS for γ-decalactone. **(F)** Violin plot showing the relative abundance of γ-decalactone in accessions carrying the MITE deletion (+, coral) versus non-carriers (-, steel blue). Horizontal lines in (-) and (+) indicate the mean.
